# Molten globule driven and self-downmodulated phase separation of a viral factory scaffold

**DOI:** 10.1101/2023.01.27.525862

**Authors:** Mariano Salgueiro, Gabriela Camporeale, Araceli Visentin, Martin Aran, Leonardo Pellizza, Sebastián Esperante, Agustín Corbat, Hernán Grecco, Belén Sousa, Ramiro Esperón, Silvia S. Borkosky, Gonzalo de Prat-Gay

## Abstract

Viral factories of liquid-like nature host transcription and replication in most viruses. The syncytial respiratory virus factories include gene function proteins, brought together by the phosphoprotein (P) RNA polymerase cofactor, present across non-segmented negative stranded RNA viruses. Homotypic liquid-liquid phase separation of RSV-P is governed by an α-helical molten globule domain, and strongly self-downmodulated by adjacent sequences. Condensation of P with the nucleoprotein N is stoichiometrically tuned, defining aggregate-droplet and droplet-dissolution boundaries. Time course analysis show small N-P nuclei gradually coalescing into large granules in transfected cells. This behavior is recapitulated in infection, with small puncta evolving to large viral factories, strongly suggesting that P-N nucleation-condensation sequentially drives viral factories. Thus, the tendency of P to undergo phase separation is moderate and latent in the full-length protein but unleashed in the presence of N or when neighboring disordered sequences are deleted. This, together with its capacity to rescue nucleoprotein-RNA aggregates suggests a role as a “solvent-protein” and possible fluidity tuner of viral factories.

## INTRODUCTION

Dynamic spatiotemporal distribution of cellular components is often achieved by the ability of macromolecules to undergo liquid-liquid phase separation (LLPS), giving rise to biomolecular condensates (BMCs) often observed as membraneless organelles (MLOs) [1-3]. In these, a variety of functions is specifically compartmentalized resulting in highly optimized biochemical processes that can be modulated by finely tuned assembly and dissolution processes. This is true for a wide spectrum of physiological and pathological reactions, and is present across life kingdoms [4-7]. The formation of BMCs often involves RNA and proteins, which require a combination of features such as: intrinsic disorder, multivalency, low complexity regions, weak interactions, modularity, oligomerization, and protein-protein and protein-nucleic acid interactions [8].

Viruses are no exception in the use of liquid-like structures to compartmentalize their gene function and particle assembly [9, 10]. These structures are referred to as viral factories and concentrate the required components for genome replication and transcription to operate, and at the same time protect and hide the genome from host innate immune mechanisms [11, 12]. Large DNA viruses of cytosolic replication were the first described to form these structures, resembling cellular aggresomes [13-15]. Single negative stranded RNA viruses (nsNSV) appear to all share the presence of cytosolic viral factories [9], with rabies virus (RABV) as a paradigmatic example. Described as Negri bodies in brain tissue in 1903, and a characteristic and diagnostic feature of rabies for decades, they were uncovered as sites for replication and transcription, with all the properties of liquid membraneless organelles [16-18]. Other examples among mononegavirales include vesicular stomatitis virus [19], ebolavirus [20], measles virus [21], borna disease virus [22], human metapneumovirus [23], and respiratory syncytial virus (RSV) [24]. Thus, since the compartmentalization of gene function through MLO-like viral factories seems to be a common feature to viruses, it calls for the investigation of the biochemical and physicochemical processes that drive and modulate their formation.

RSV is an enveloped non-segmented negative-stranded (nsNSV) RNA virus that belongs to the pneumoviridae family, is an archetypical member of the Mononegavirales order, and a model system to address the underlying mechanisms of viral factory assembly. The replication complexes of these viruses comprise an RNA-dependent RNA polymerase (L), a phosphoprotein (P) and a nucleoprotein (N) packed with genomic and anti-genomic RNA. Transcription starts at the 3’ leader region of the 15 kb genome and proceeds through “gene start” (GS) and “gene end” (GE) sequences, yielding 10 individual transcripts that code for 11 proteins [25-27]. RSV include a unique transcriptional antiterminator, M_2-1_, which ensures maximum processivity of transcription towards the 5’ end of the polar genome [28]. In fact, the RNA polymerase complex of RSV was shown to condense into viral factories within infected cells, with a subcompartment where M_2-1_ and newly synthesized viral mRNA, and not the other components (N, L, and P), accumulate [24].

Cytoplasmic inclusion bodies were observed when RSV N and P were co-transfected [29], something subsequently observed in related viruses [17, 21, 30, 31] indicating both proteins as minimal requirements for the formation of inclusions *in vivo* and *in vitro* [32, 33]. As in most nsNSVs Mononegavirales, the nucleocapsid of RSV is characterized by rod-shaped structures of helical symmetry with ten contiguous N protomers per turn forming a continuous groove that accommodates genomic RNA between the N- and C-terminal domains of the 391aa N protein (Figure 1A) [34, 35]. Similar to other Mononegavirales, recombinant expression of RSV N yields decameric ring-like structures wrapped by 70 nt long RNA molecules from the host cell (N_RNA_ rings), where removal of RNA can only be achieved under strong chemical denaturation conditions and leads to irreversible aggregation of RNA free N (Figure 1B) [36-39].

**Figure 1.**
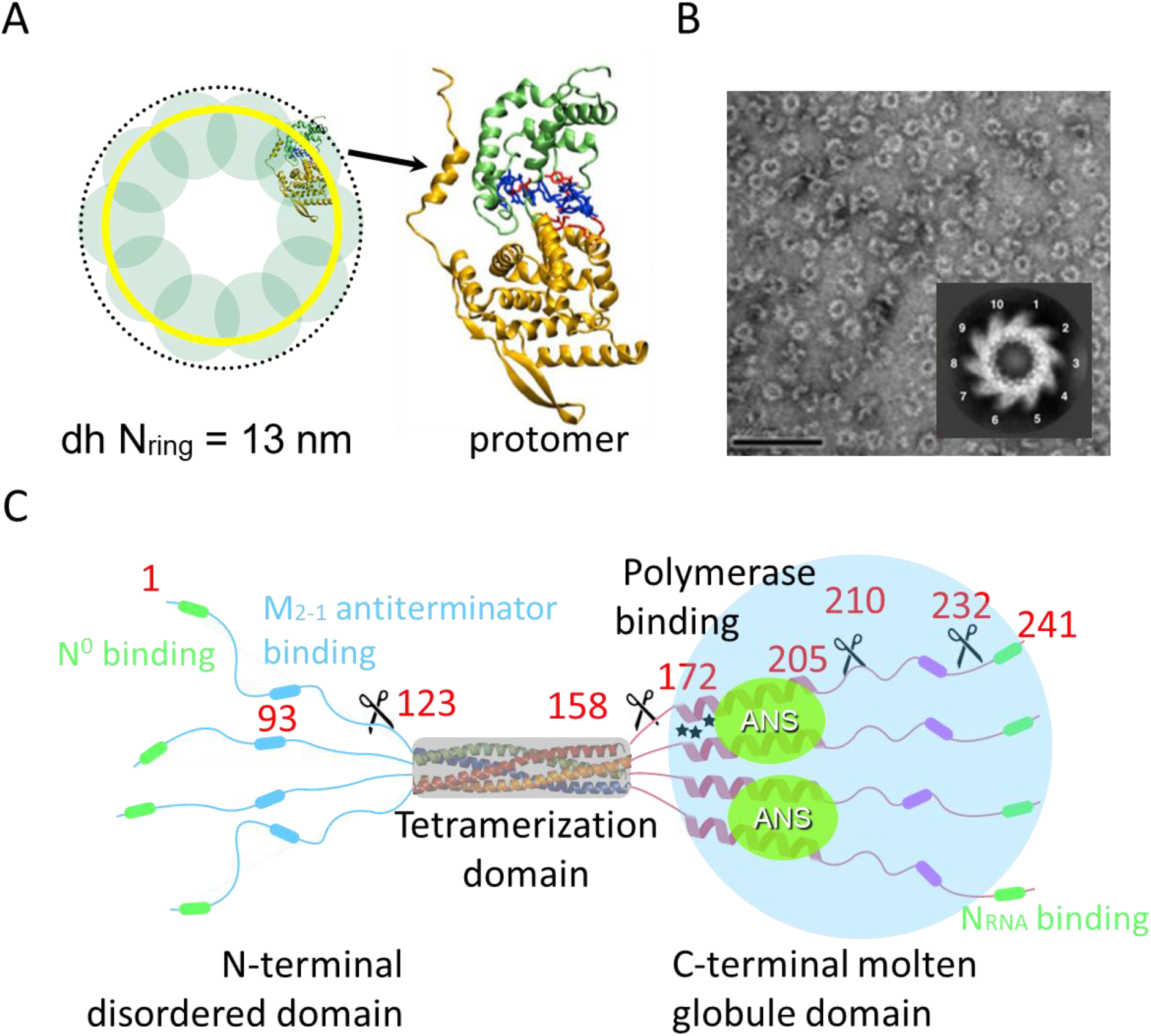
Structure and conformational properties of RSV nucleocapsid N and phosphoprotein P. **(A)** N wraps the viral genome forming a helical nucleocapsid and recombinant expression yields NRNA rings mainly composed of 10 protomers corresponding to one turn of the nucleocapsid, strongly bound to host non-specific RNA. **(B)** Transmission electron microscopy images of 13 nm diameter NRNA rings. **(C)** RSV-P is a tetramer of 15 nm diameter composed of a highly stable, four helix bundle tetramerization domain flanked by an N-terminal intrinsically disordered domain and a C-terminal molten globule, ANS binding domain 50. It contains linear binding motifs for N_0_, N_RNA_, and M_2-1_, and an L-binding surface that spans the tetramerization domain and the C-terminal adjacent region.

Initially described as a polymerase cofactor, the phosphoprotein P operates at the center of transcription and replication in Mononegavirales and binds N, L and, in the case of RSV and pneumoviruses, M_2-1_ [40-49]. The phosphoproteins alternate disordered regions with linear binding motifs with tight α-helical oligomerization domains, and although they are completely divergent in sequences, they share function and domain arrangement, in particular those of the Pneumoviridae family [50]. Their flexible domains have different degrees of disorder and we experimentally showed that while the N-terminal domain of RSV-P is an intrinsically disordered domain (IDD) as predicted, the C-terminal domain is an α-helix rich molten globule (αMG) with a low stability but cooperative folding module [41, 51] (Figure 1C). This was subsequently confirmed by NMR chemical shift analyses and α-helix propensities of the isolated C-terminal fragment monomer since this region is not detectable at room temperature in the context of the full-length P tetramer [52, 53]. Only α-helix propensities and not persistent structure can be observed in the monomeric domain fragment, the αMG only forms in the context of the tetramer, indicating interprotomer interactions favored by tethering to the four-helix bundle tight oligomerization domain [51].

In this work, we show that homotypic phase separation of P is driven by the stability of a α-helical molten globule and is strongly downmodulated by adjacent sequence regions. We show that the heterotypic condensates formed with the N_RNA_ nucleocapsid model are finely tuned across aggregate-droplet-dissolution boundaries, according to P concentration. These condensates start in the cell from nuclei formed by recently synthesized P and N and gradually coalesce into a discrete number of large condensates, a phenomenon that may be recapitulated in virus infection. The established liquid nature of RSV’s viral factories [24, 32], recent work on HMPV phosphoprotein [54], and the fact that P is at the center of the interaction network within the replication complex, in combination with our results suggest P as the scaffold that not only drives the viral condensation but may tune the fluidity and thus play a crucial role in viral factory formation and fate.

## RESULTS

### RSV-P undergoes homotypic LLPS that is strongly modulated by adjacent sequences at the C-terminus

RSV P displays the paradigmatic features of an LLPS scaffold: intrinsically disordered regions (IDRs), multivalency, stable oligomerization, modular architecture, high stability, and both weak and strong interactions with other viral and cellular proteins. As we increased both protein and crowder concentrations (see materials and methods), typical spherical homotypic droplets were observed by microscopy (Figure 2A, 2B, and S1A). Protein concentration inside droplets reached 6.7 mM compared to 7.1 µM in the dilute phase (Table S1). Homotypic LLPS is maximum at low temperature as revealed by turbidity, it is dissolved at 45°C, and returns upon lowering the temperature back to 15 °C, indicating full reversibility (Figure 2B, right). Droplet coalescence was observed (Figure 2C), and dynamic behavior was further supported by the immediate incorporation and homogeneous distribution of fluorescently labelled P (P*) into pre-formed unlabeled P-droplets (Figure 2D), incorporation and therefore dynamic nature persists after a 4 h incubation (not shown). Fluorescence recovery (FRAP) experiments show recoveries around 50% and a k_obs_ of 8.8 ± 1.7 10^−3^ s^-1^ corresponding to a t_1/2_ of 80 seconds (Figure 2E).

**Figure 2.**
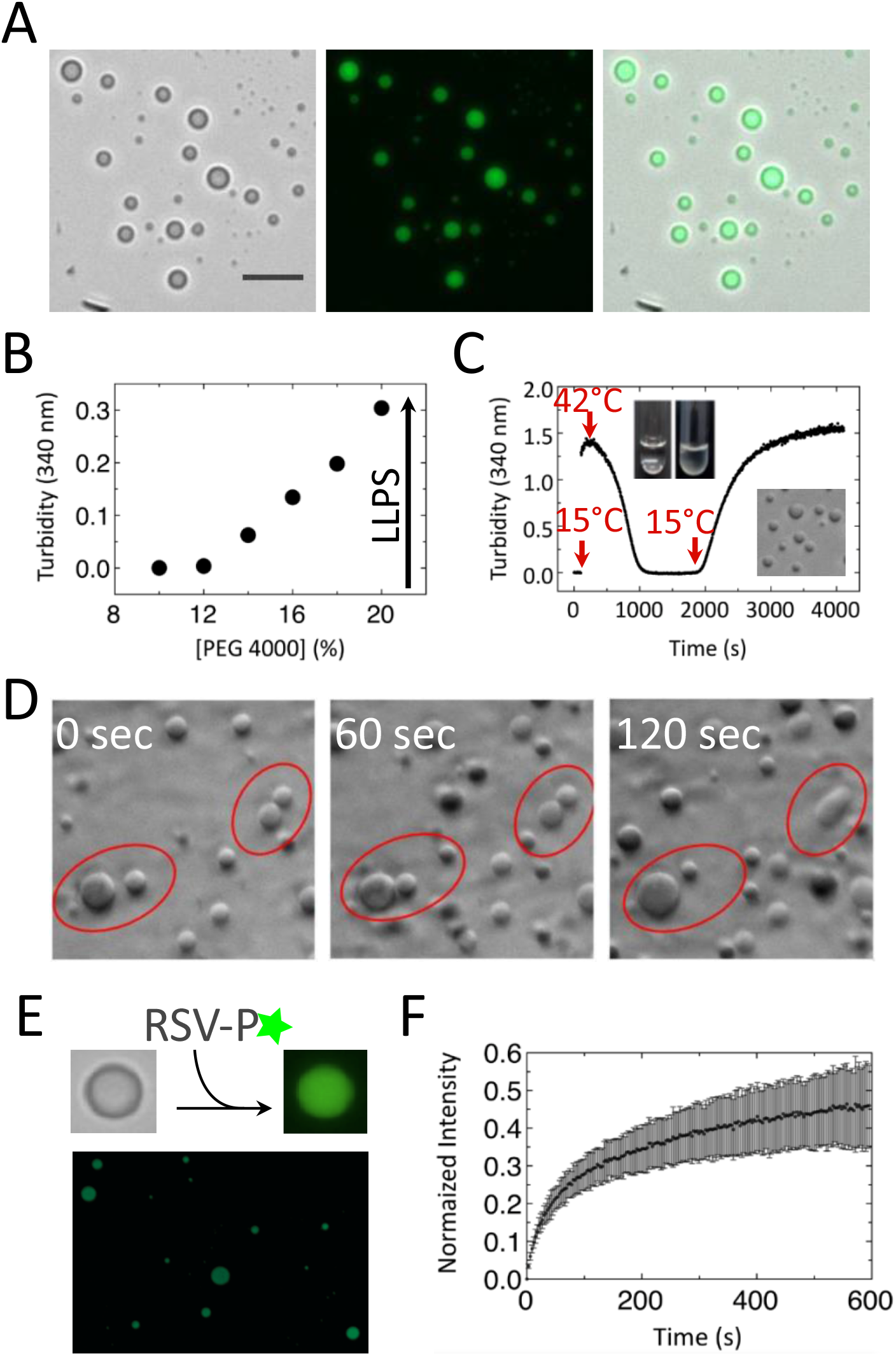
Homotypic LLPS of RSV-P. **(A)** Brightfield and fluorescence microscopy images at 40X of 25 µM P (125 nM FITC-P). Scale bar = 30 µm. **(B)** Effect of crowding on homotypic LLPS monitored by turbidity. 5 µM P was incubated in the presence of increasing concentrations of PEG 4000 and the onset of LLPS was evaluated by measuring turbidity at 340 nm. **(C)** Effect of temperature on homotypic LLPS. 2.5 µM P was added to reference homotypic buffer at 10°C and droplet formation/dissolution was monitored by measuring absorbance at 340 nm as the sample was heated to 45°C, and then cooled back to 10°C. Initiation of temperature changes are indicated with red arrows. Images of P samples at low and high temperatures are shown as an inset to denote the difference in turbidity, and microscopy images show droplets as the temperature is returned to 15°C. **(D)** Coalescent events of homotypic droplets registered by bright field microscopy at 40X magnification at different times. **(E)** Incorporation and distribution of FITC-labelled P to preformed homotypic non-labelled droplets was determined by fluorescence microscopy. **(F)** Quantitative analysis of homotypic LLPS dynamics by FRAP (materials and methods). Values represent mean ± s.d.

In order to address domain structure determinants of the homotypic LLPS process, we constructed a series of variants (Figure 3A): *i)* PTetC, covering the tetramerization and C-terminal domains, *ii)* PNTet, corresponding to N-terminal and tetramerization domains, and *iii)* two C-terminal deletions that lack the linear binding motif to the nucleocapsid (P_Δ232_) and the adjacent unstructured region (P_Δ210_) [53]. Except for PTetC, all constructs yield droplets in the reference condition (Figure 3B, see Methods) but PNTet condensates exhibit irregular shape, which implies that both N- and C-domains are involved in the demixing process, and that the C-domain modulates the material properties of the demixed state, possibly altering the interaction dynamics that drive LLPS, or other properties. For a quantitative analysis of the tendency of the variants to demix, we carried out thermal scans followed by turbidity and determined the threshold temperature (T_LLPS_) at which each species starts demixing, with the full-length wild-type P starting at 33°C (Figure 3C). PTetC does not demix even at temperatures below 10°C, indicating that the N-terminal domain is required in this concentration range. However, at higher crowder and protein concentrations, PTetC can form reversible droplets that exhibit similar behavior towards temperature and ionic strength as full-length wild-type P (Not shown). PNTet tendency to phase separate is much lower than that of full-length P (T_LLPS_ = 20°C), further supporting the crucial role of C-terminal domain to promote LLPS. Interestingly, the LLPS tendency of the deletion variants to yield regular droplets increased dramatically, with T_LLPS_ of 44°C for the P_Δ232_ and 55°C for the P_Δ210_ variants, respectively. Clearly, there is sequence-intrinsic disorder information within the deleted regions that negatively and strongly modulate the homotypic LLPS of P (all transitions were fully reversible, Figure S2A). As in the case of the full-length P, all variants that form droplets are able to immediately recruit labelled-protein from the diluted phase, indicative of a dynamic nature (Figure S2B).

**Figure 3.**
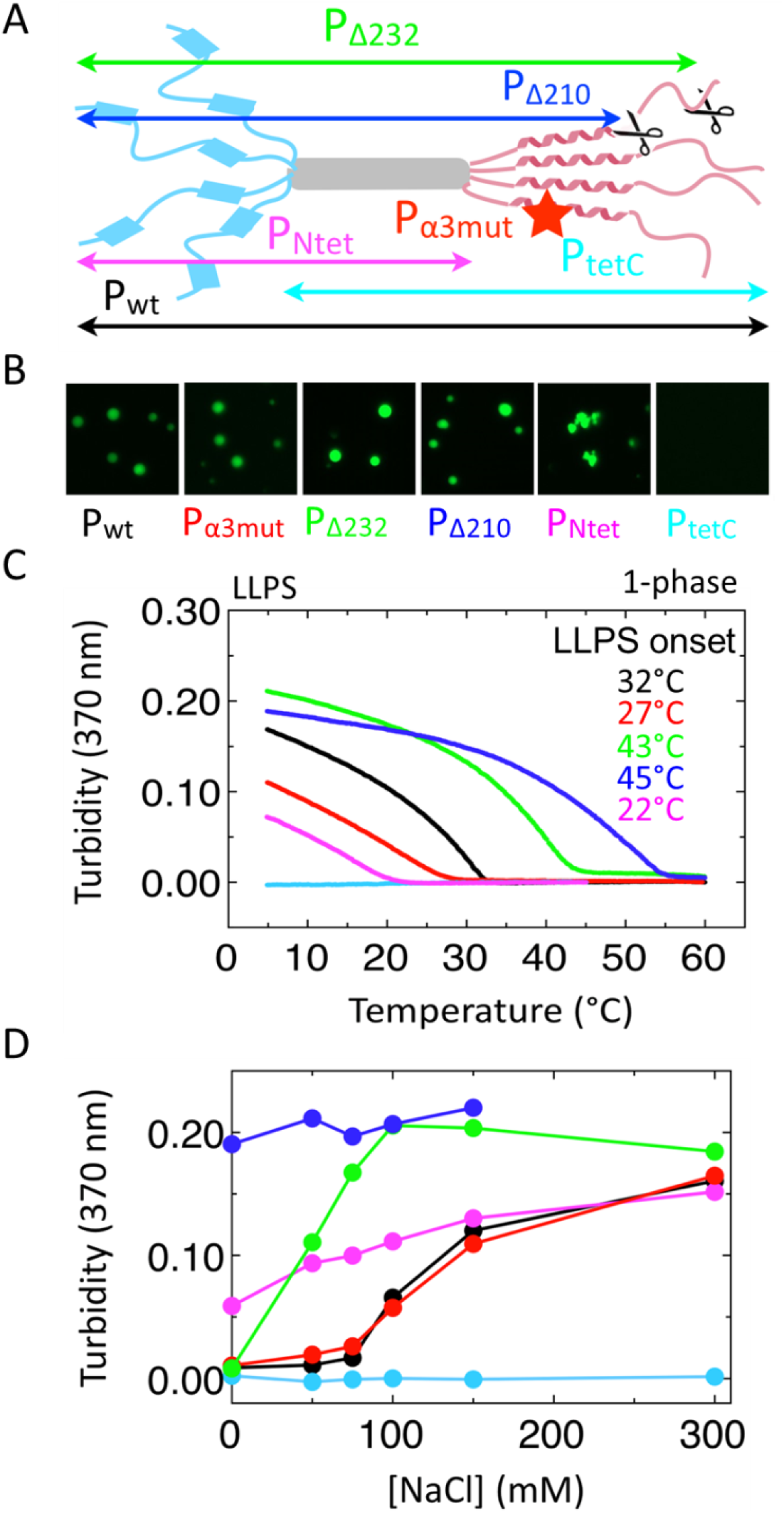
Deletion and quantitative analysis of homotypic RSV-P LLPS. (**A)** Description of P constructs: WT (P full length), PTetC (P_103-241_), PNTet (P_1-160_), P_Δ232_ (P_1-231_), P_Δ210_ (P_1-209_), α3mut (P full length with substitutions G172S, R174A and E175G, indicated with a star at the N-cap of α-C1). **(B)** Analysis of demixing of P constructs by fluorescence microscopy at 40X. All samples contain 12.5 µM protein and were incubated for 1 h at 20°C before imaging. Scale bar = 20 µm. **(C)** Comparison of LLPS propensities between P constructs as a function of temperature. Threshold temperatures for protein demixing were determined as the onset of turbidity. 5 µM protein was incubated at 65°C (WT, black; α3mut, red; Δ210, blue; Δ232, green) or 45°C (PTetC, cyan; and PNTet, magenta) for ten minutes and turbidity at 370 nm was monitored as the sample was cooled down to 5°C at a rate of 1°C/min. **(D)** Comparison of LLPS propensities between P constructs as a function of NaCl concentration. 12.5 µM protein was incubated for 1 h at 20°C and turbidity was measured at 370 nm.

The demixing phenomenon is very sensitive to ionic strength, being impaired at low salt concentrations as shown both by fluorescence microscopy and turbidity measurements (Figure S53A). Although droplets tend to deform at high ionic strength (> 200 mM), frequent coalescence events can be observed over a wide range of salt concentrations, indicative of a liquid-like state (Figure S3B). By measuring turbidity as a function of salt concentration for all P variants, we found that total or partial deletion of the C-terminal domain affects LLPS sensitivity to ionic strength (Figure 3D). P_Δ232_ turbidity reaches a plateau at lower NaCl concentration than full length P. Furthermore, P_Δ210_ turbidity signal is completely insensitive to ionic strength, with demixing taking place even in the absence of salt. This suggests that the putative negative regulatory element located among the last 30 residues of P might exert its effect through electrostatic repulsion. Thus, a salting-in effect could explain why low NaCl concentrations are required for full length P demixing, consistent with strong contribution by hydrophobic interactions. In agreement with the temperature scans, this element has more than one component, as there is a partial and differential effect between the two deletions. PNTet shows a mild dependence with the ionic strength, supporting that although it has an impact on homotypic LLPS, it is not a main determinant as it has low charge. PTetC shows no demixing across the ionic strength range analyzed at this protein concentration.

### The stability of the α-helical molten globule at the C-terminal domain tunes homotypic LLPS

As we previously described, the C-terminal domain has molten globule (MG)-like properties in the context of the tetramer, namely, maximum stability at low temperature, it unfolds completely by 45 °C, and is independent of the intrinsically disordered N-terminal domain [51]. The temperature-induced unfolding transition of this domain is accompanied by a CD change in parallel with the homotypic LLPS, as indicated by the superposition with the turbidity signal (Figure 4A). These results, together with the full reversibility of both processes are indicative of a strong correlation between conformational stability of the C-terminal αMG domain of P and its tendency to demix, thus supporting a role for this structure in mediating homotypic intermolecular interactions that promote LLPS. In this context, conditions that stabilize these α-helical structures would show an increased tendency to undergo LLPS, where destabilization of the structure would have the opposite effect. No significative effect of either NaCl or PEG was observed on the MG thermal unfolding.

**Figure 4.**
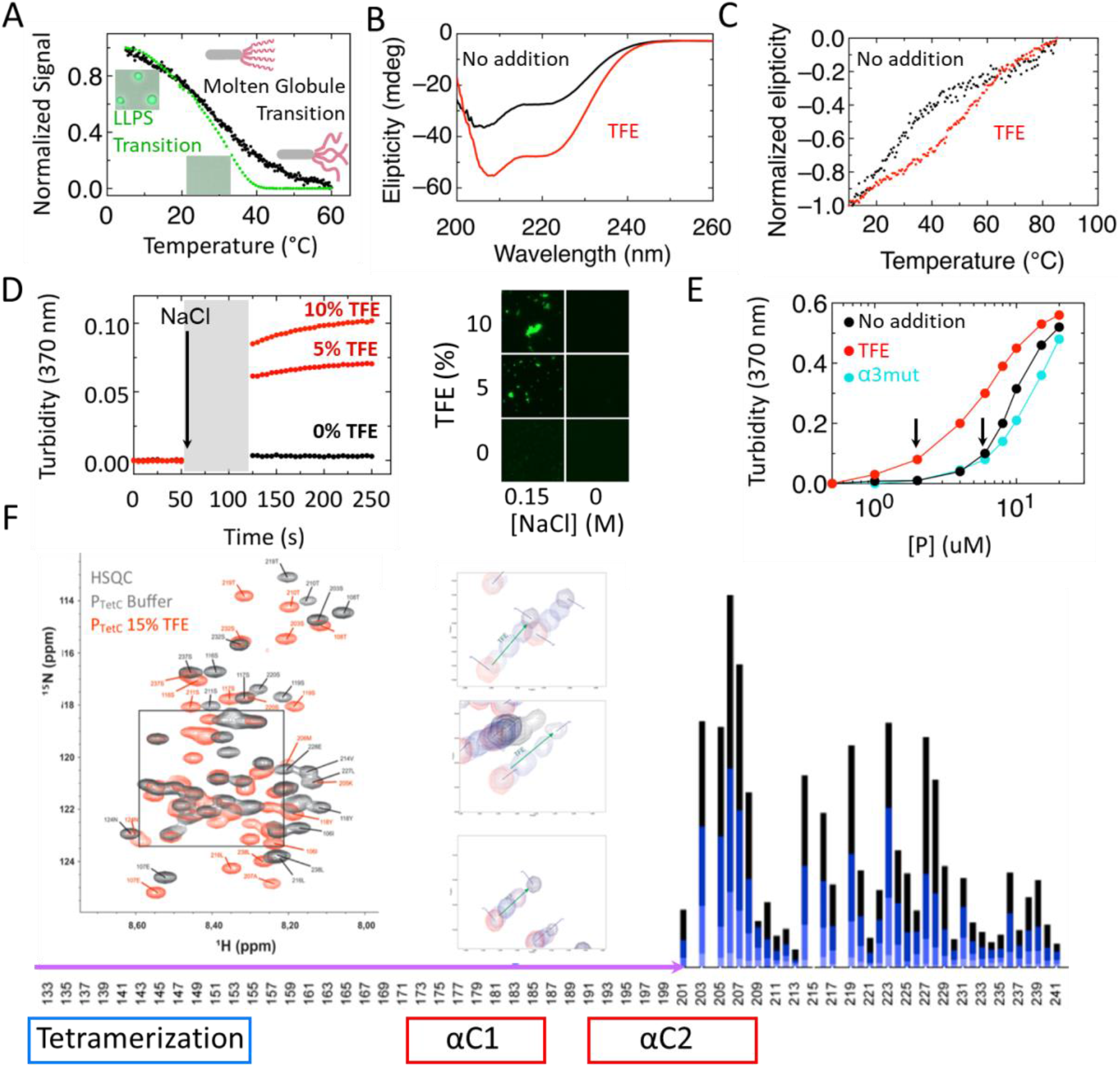
Stability of the α-helical MG domain (αMG) of P drives homotypic LLPS. **(A)** Temperature-induced P homotypic droplet dissolution followed by turbidity (green line) and αMG destabilization followed by far UV CD at 220 nm (black line). Turbidity experiment (5 µM P WT in reference buffer), and for the CD unfolding experiment, 3.75 µM P WT in 10 mM Tris-HCl, 25 mM NaCl, pH 8). **(B)** Far UV CD spectra of P WT (black) and with 20% TFE added (red). **(C)** Thermal scans followed by CD at 220 nm of P WT (black) and with 20% TFE added (red). **(D)** Effect of TFE on P WT condensation. Demixing of 2.5 µM P in reference buffer was evaluated by turbidity at 370 nm after addition of 0.15 M NaCl in the absence (black circles) and presence (red circles) of 5% and 10% TFE, respectively. Fluorescein-labelled P samples were then prepared in the absence and presence of 0.15 M NaCl and analyzed by fluorescence microscopy at 20°C. Scale bar = 20 µm. **(E)** LLPS tendency of P WT (black), P WT plus 5% TFE (red), and α3mut (cyan) were determined by dilution in reference buffer at 15°C. Arrows indicate critical concentrations for P WT in 5% TFE (1 µM) and P WT (4 µM) and α3mut (5 µM). **(F)** HSQC spectra of PTetC (black) and with 15% TFE added (red). Three selected peaks representative of gradual titration with TFE (green arrow). Addition of TFE from light blue to black represent the degree of chemical shift variation for each residue affected. Regions spanned by part of the tetramerization domain (blue) and αC1 and αC2 (red) are shown in boxes. Only residues from 133 onwards are represented for clarity. Arrow in magenta indicates the NMR invisible region.

The difficulty in extracting structural details precludes modeling mutations that may increase the stability of the αC1 helix within the αMG, something that would be difficult even with an atomic structure at hand. In order to address this, we made use of solvent stabilization. We had previously shown that trifluoroethanol (TFE) stabilizes the pre-formed α-helical content in the C-terminal domain which can be accurately estimated from the CD spectra to be 32 residues [51] subsequently confirmed by α-helical propensity from NMR chemical shift analysis [53]. Addition of 20% TFE shows a marked increase in α-helical content as judged by the far UV CD spectrum (Figure 4B). TFE titrations of both P and PTetC show a superimposable change in secondary structure in agreement with the independence from the N-terminal domain, with a plateau at 20% (Figure S4A). Moreover, 20% TFE stabilizes the domain towards thermal unfolding by an 11°C increase in T_m_ (Figure 4C). In order to test the correlation between helix stabilization and tendency to LLPS, we analyzed the effect of TFE on the demixing process. As little as 5% TFE largely favors homotypic LLPS, as indicated by the change in turbidity and the formation of condensates by fluorescence microscopy, at experimental conditions that would otherwise not allow demixing (Figure 4D). As a control, condensation does not occur in the absence of salt, indicating that the same LLPS driving forces continue operating in the presence of TFE, and that TFE alone cannot induce LLPS (Figure S4B). A quantitative picture of this effect is evidenced by a 4-fold decrease in the critical protein concentration required for LLPS even at 5 % TFE (Figure 4E). We next carried out an NMR analysis by performing HSQC spectra in the absence or presence of 15% TFE (Figure 4F). As described [53], most of the αMG region shows no NMR signals but we observe a large stabilization of the C-terminal half of αC2, as indicated by the large change in chemical shifts of contiguous residues.

In the absence of a defined atomic structure and given the undetectability of the NMR signals in the region that flanks the tetramerization domain (Figure 1C), we used the helical propensities to define the N-cap of αC1 at positions conserved across pneumoviruses [53]. Thus, in order to generate a α-helix destabilizing mutation, we produced a triple mutant of three conserved residues (G172S R174A E175G), and name it α3mut. The effect of this mutant on helix stability of the αMG is confirmed by the disappearance of the unfolding CD transition between 10 and 45°C (Figure S4C). The hypothesis was further confirmed by a substantially lower tendency for LLPS observed in quantitative cooling transition analysis (Figure 3C), supporting a role for αC1 helix as a driving force in the homotypic LLPS. The mutation also shows a moderate but significantly higher critical protein concentration for LLPS (Figure 4E).

### P and N_RNA_ rings undergo stoichiometrically tuned heterotypic LLPS

Heterotypic condensation into regular spherical droplets was readily observed after mixing P and N_RNA_ at protein and crowder concentrations much lower than those required for homotypic phase separation of P and it was also observed even in the absence of crowder (Figure 5A and S5). All droplets contain both proteins and their average size increases with concentration in excess of either of the components (Figure 5B) or at increasing both components at fixed ratios (Fig S6A, diagonals). The formation and dissolution of the droplets is governed by the relative concentration of the components at a stoichiometric boundary under excess of P, from 5P : 1N_RNA_ to 10P : 1N_RNA_ (Figure 5B and S6B). Dissolution of the NP condensates by excess of P follows the concept of re-entrant phase after saturating the N binding sites and breaking the interaction network [55]. A transition boundary between ordered aggregates and droplets is also observed in excess of N_RNA_ from 2.5P : 4N_RNA_ to 1.25P : 4N_RNA_ (Figure S6B). When we incubate N_RNA_ alone we observe an aggregate that is immediately and completely rescued into condensates by addition of P (Figure 5C). Altogether, these results indicate a determinant role for P in tuning the condensate’s fluidity and N_RNA_ solubilization and recruitment into the condensates. Using hydrophobic coating for the coverslip, we observe that the heterotypic N_RNA_-P droplets wet the glass, while the homotypic P do not, highlighting differences in surface properties (Figure 5D), which suggest a hydrophobic nature for the N-P condensate’s surface.

**Figure 5.**
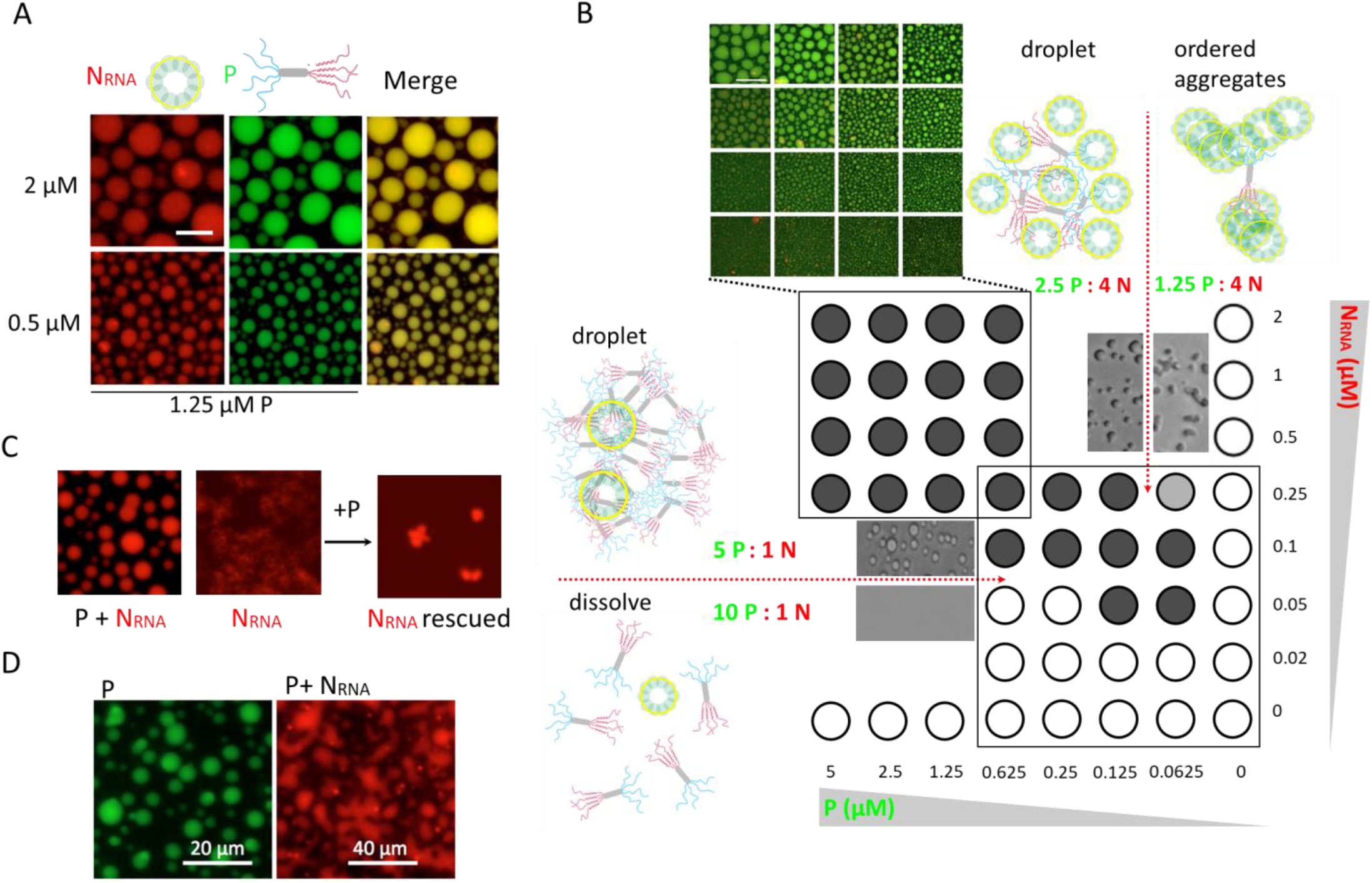
Heterotypic P and N LLPS. **(A)** Colocalization of Cy5-labeled N_RNA_ and Fluorescein-labeled P WT into demixed droplets determined by fluorescence microscopy at two molar ratios (Scale bar = 10 µm). **(B)** Two heterotypic LLPS grids spanning different concentrations ranges of N_RNA_ and P WT are schematically representing the demixed state as black full circles, homogeneous mixed state or amorphous aggregation as empty circles, and a gray full circle is used to indicate a singular non-spherical demixing condition. Red dashed arrows indicate two distinct phase transition events: (i) demixed droplets to homogeneous solution and (ii) demixed droplets to ordered aggregates. Light microscopy images and molar ratios from the mixtures involved in those phase transitions are shown, along with model representations. **(C)** Fluorescence microscopy images of samples containing P WT + Cy5-labeled N_RNA_ in the presence of 5% PEG 4000. Addition of P WT after incubation of N_RNA_ alone for 1 h rescues N_RNA_ from an aggregated state to a condensed state. **(D)** Demixed samples of P or P+N_RNA_ were loaded between two coverslips pre-treated with a hydrophobic coating (Sigmacote®) and analyzed by fluorescence microscopy at 40X. Homotypic sample: 12.5 µM P. Heterotypic sample: 5 µM P, 0.5 µM N_RNA_.

Next, we wanted to assess the role of domains and regions of P, using the same set of constructs used for probing homotypic LLPS (Figure 3A). At a ratio of 5P: 4N_RNA_, full-length wild-type P and the α3mut mutant form indistinguishable droplets, and PTetC from regular but smaller droplets (Figure 6A). The other species, PNTet, P_Δ210_, and P_Δ232_ do not form droplets but yield co-aggregates that containing both P and N_RNA_, suggestive they are product of interaction but not productive LLPS. A closer quantitative analysis of different ratios of mixtures (Figure 6B and S7) shows that increasing the concentration of PNTet yield smaller albeit regular droplets from 2.5PNTet : 1N_RNA_ ratio upwards, indicating that interactions with the nucleoprotein leading to LLPS can take place without the C-terminal domain of P under these conditions. The fact that P_Δ210_ and P_Δ232_ form co-aggregates with N_RNA_ at intermediate ratios, is also in line with the idea that interactions with N_RNA_ can also occur even in the absence of the binding site previously mapped to the last 10 residues of the C-terminal domain of P [48] and secondary interaction sites were proposed by paramagnetic NMR [53].

**Figure 6.**
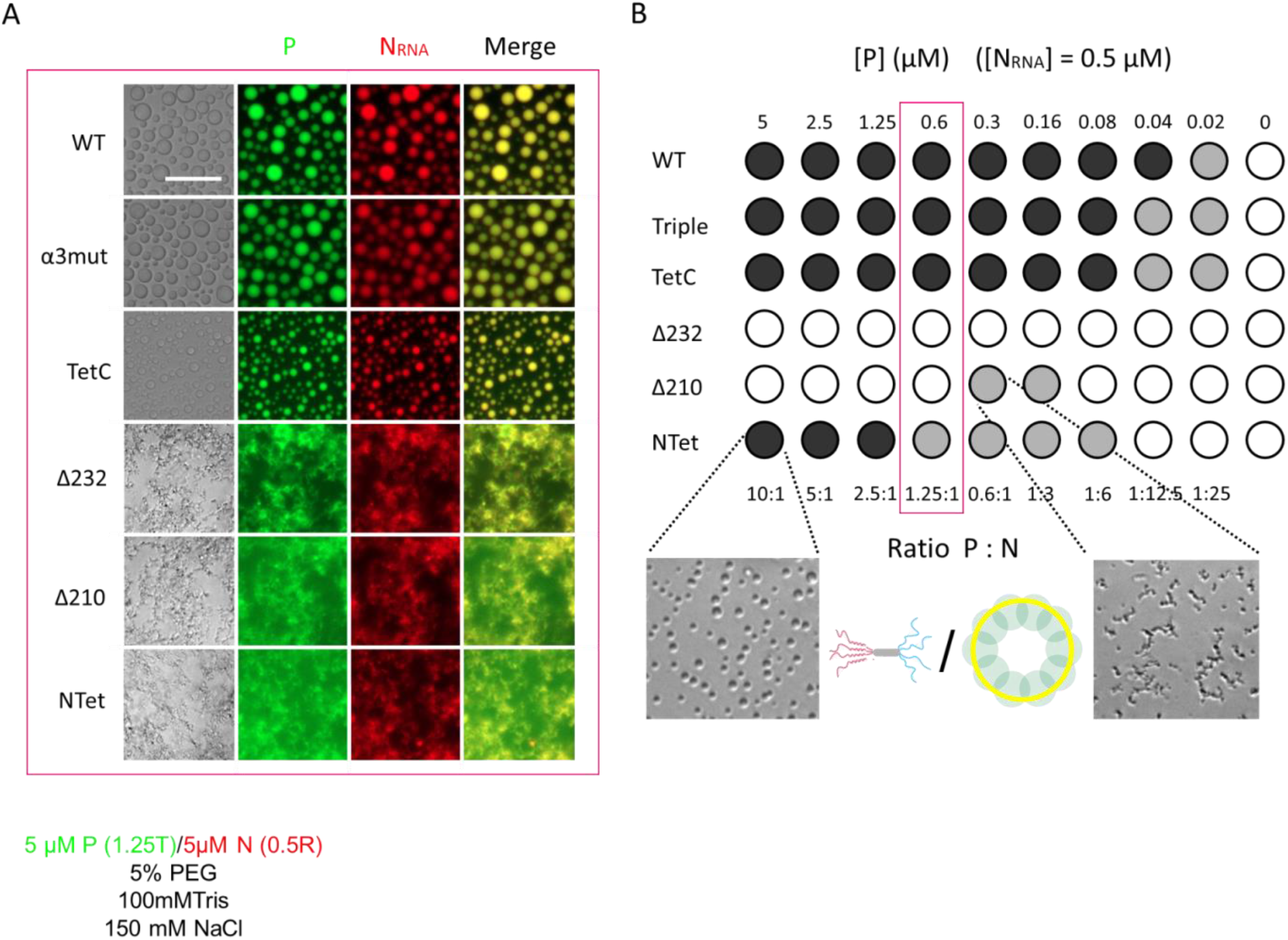
Domain and P deletion analysis of heterotypic P-N LLPS. All P variants were tested for its capacity to form heterotypic droplets with N_RNA_ at different molar ratios, spanning a wide range of P concentrations while keeping [N_RNA_] fixed at 0.5 µM. **(A)** A single molar ratio was selected (red rectangle in the figure 6B grid) to show colocalization of P variants and N either into spherical droplets or amorphous aggregates. **(B)** A schematical representation of the grid is shown: droplets (black full circles), non-regular condensates (gray full circles), amorphous aggregates or no condensation (empty circles), see Figure S7 for details. P concentrations are shown on the upper side of the grid and molar ratios P: N_RNA_, on the lower side. Images showing PNTet LLPS at 10:1 P:N_RNA_ ratio and P_Δ210_ LLPS at 0.6:1 P:N_RNA_ ratio are displayed.

### Dynamic P-N_RNA_ condensates in the cell as drivers of viral factory assembly

As the next step in the analysis of the heterotypic P-N_RNA_ LLPS, we investigated their occurrence and nature within cells. For this, A549 cells were transfected with a GFP fusion of P and unfused N, and analyzed by fluorescence microscopy. While both proteins show a diffuse pattern when expressed separately, co-expression shows that P and N constitute the minimum requirement for the formation of the granules [29, 32, 44]. These are visualized as 1-4 µm in diameter bodies distributed mostly at the perinuclear region, equally detectable by GFP-P fluorescence and immunostaining of both N and P, indicating that the GFP fusion at the N-terminus of P does not affect the process (Figure 7A and 7B, left). In addition, we observe that PTetC is capable of forming granules indistinguishable from P, most likely because of the high concentrations within the intracellular granules or the presence of other yet undefined cellular components (Figure 7B, center). PNTet does not form granules, as oppose of what is observed *in vitro* (Figure 7B, right). In order to determine whether these granules represent liquid-like entities we investigated their dynamics using fluorescence recovery after photobleaching (FRAP). An almost 100% fluorescence recovery of the granules takes place after 2 seconds, indicating a highly dynamic process and a liquid-like nature of the condensate (Figure 7C). Next, we analyzed the time course formation for the in cell P-N condensates. For this, we transfected A549 cells with the same GFP-P and N containing vectors, and observed the evolution of the process (Figure 7D, Movie S1). At 9 h post-transfection, a large number of small puncta are observed which evolve to a discrete number of granules of much larger size up to 15 hours post transfection. This result strongly suggests that coalescence of P-N nucleus formed by newly synthesized proteins that evolve into large condensates drives the formation of viral factories. We infected A549 cells and immunostained for both P and N proteins at different time points. At 24 h post-infection we observe the viral factories assembled [24] (Figure 7E, arrows), and at intermediate times we observe small size granules that grow as the infection progresses (Figure 7E, P immunostaining shown).

**Figure 7.**
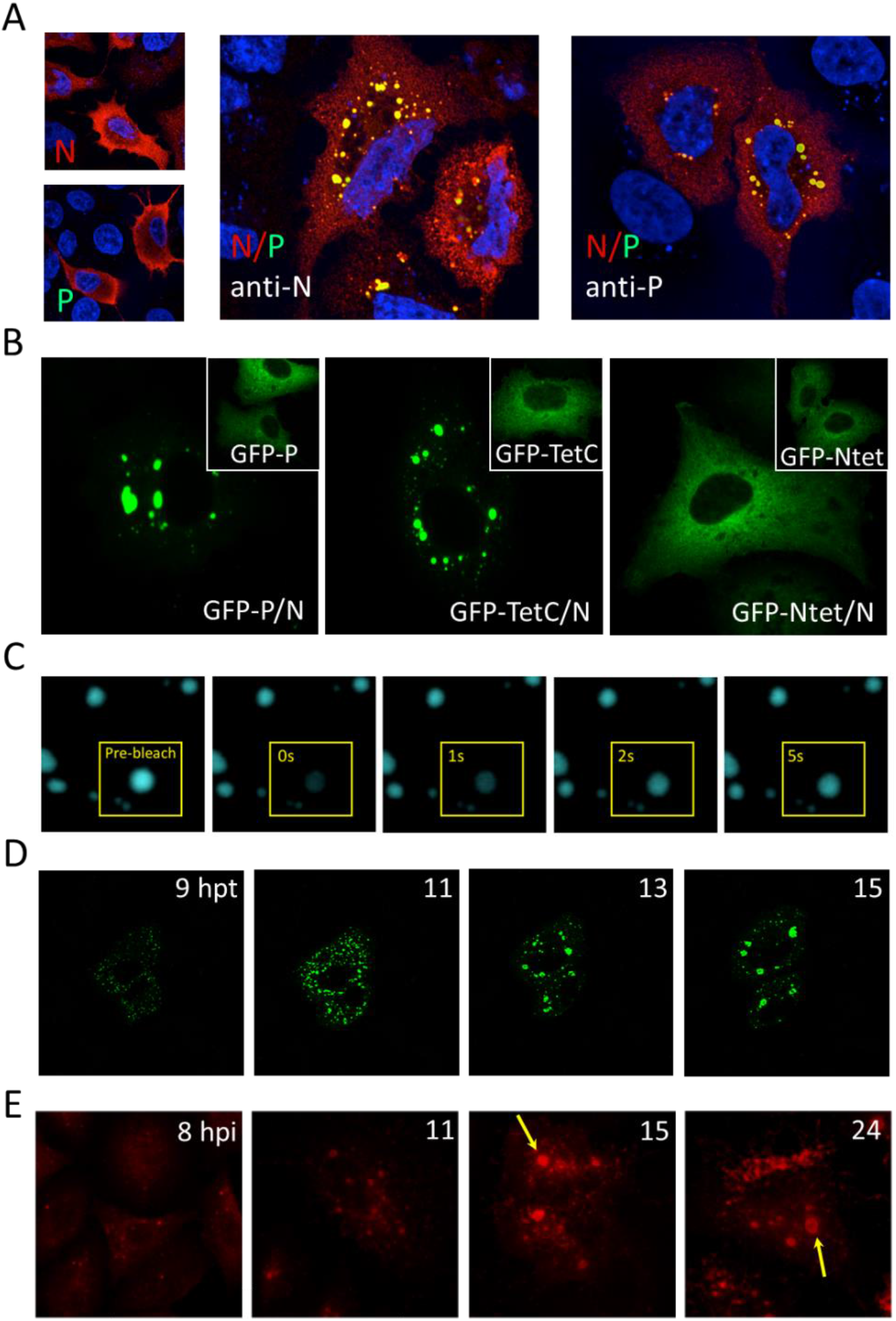
P-N_RNA_ condensates as drivers of viral factory assembly in A549 cells. **(A)** Viral protein distribution in cells transfected with pCDNA-N or –P alone (left) or co-transfected with both plasmids and detected by fluorescence microscopy labeling with anti-P or anti-N antibodies. Nuclei were stained with DAPI. **(B)** Viral protein distribution in cells co-transfected with pCDNA-N and pCDNA-GFP.P (left), - GFP.TetC (center), or –GFP.NTet (right). The insets show the diffuse distribution in the cytoplasm of GFP-P, GFP-PTetC and GFP-PNTet expressed alone. Viral proteins were observed by fluorescence microscopy at 488nm. **(C)** FRAP of condensate after 24h co-transfection with pcDNA-N and pCDNA-GFP.P plasmids. **(D)** Time course formation of P-N condensates in A549 cells co-transfected with pcDNA-N and pCDNA-GFP.P plasmids. The evolution of the granules was registered using a Zeiss LSM 880 Airyscan confocal laser-scanning microscope at 37°C and 5%CO_2_ over 15 hpt. **(E)** P immunostaining at different time points of A549 cells infected with RSV-A2 strain (MOI=1). Viral factories at 15 and 24 hpi are pointed (yellow arrows).

## DISCUSSION

Phosphoproteins from non-segmented mononegavirales are polymerase cofactors that share functional and domain arrangements as well as structural features, without sequence similarity. They also share the capacity to interact with all other members of the replication complex, and with matrix proteins. These events raise them as hubs or scaffolds for viral protein activities around replication. In this paper we first show how RSV P can undergo fully reversible homotypic LLPS favored by low temperature and optimal ionic strength with a partition coefficient ca. 1,000, roughly 200 mg/mL within the droplet. This process is dynamic as indicated by immediate incorporation of added protein to the droplets, coalescence, and FRAP experiments. Recent work showed that HMPV P has more tendency to LLPS compared to RSV P, in the absence of crowder at high protein concentrations (5 mg/mL) [54]. The RABV P was also shown to undergo homotypic LLPS [56].

We have previously determined that the C-terminal domain of P includes the thermosensitive αMG structural element which helical propensity was subsequently mapped to two α-helices α_C1_ and α_C2_ spanning residues 172-206 in the isolated monomeric domain (Figure 4F). However, no signals for this region and the oligomerization domain are observable by NMR in the tetrameric P, most likely due to dynamic conformational changes at the nanosecond-millisecond timescale [53]. In this work, we confirm that the structural αMG element consists of these two NMR invisible helices since deletion of residues 210 to 241 still show the full secondary structure and thermal transition by CD, and these helices are stabilized by being tethered to the tetramerization domain, strongly implying they are stabilized by interprotomeric, i.e. quaternary, dynamic interactions. In excellent agreement with our results, recent studies show that at 40°C the “invisible” signals appear, consistent with unfolding of the αMG [52].

Interestingly, the recent structure of the polymerase L-P complex showed an unusual tentacular arrangement comprised of a three helix bundle, in which one of the three protomers contacts the polymerase extensively [57, 58]. This helix spans residues 174-185, coincident with α_C1_, highly conserved among pneumoviruses and part of the αMG. In addition, the fourth protomer makes extensive contacts with other regions of the polymerase, showing an overall chameleonic structure and recognition capacities that nevertheless lead to a tight interaction that stabilizes the RNA polymerase. This confirms an evolutionary conserved structural plasticity of this region, clearly far from being coil or intrinsically disordered, unlike the disordered regions located towards de C-terminus of P (210-241). The ability to interact with very different surfaces on the polymerase highlight the role of the tetramerization domain in stabilizing a collection of several otherwise weak interactions that would not recognize the polymerase as monomers.

A key finding in this work is the link between the stability of αMG and the tendency to undergo homotypic LLPS, where low temperature stabilizes the αMG and LLPS, and the opposite is true at ca 40°C. We probed this hypothesis using solvent stabilization of α-helix by TFE, which both stabilize the αMG and increases LLPS tendency. Helical propensity was described to increase LLPS in TDp43 [59, 60], but no stable secondary structure as the one in the P αMG. For probing the effect of helix destabilization on LLPS by mutation, we used triple mutant on the N-cap of the transient α-helix C1 (α3mut), designed to destabilize this structure and establish its role in LLPS. Natural single residue substitutions in these amino acid positions were shown to affect viral proliferation, reducing the optimal temperature for virion production from 37 to 30°C [61]. One possible explanation is proposed that its effect would arise from weakening contacts between P and L [58]. An alternative and not mutually exclusive explanation is that the change in optimal growth temperature could be a reflection of an impaired condensation tendency of the viral factories. We must consider that P is in large excess over L in infected cells and is the scaffold for the formation of the heterotypic condensates likely to drive the formation of the viral factories. Therefore, P-L contacts being affected would not be the unique source of mutation effect of the αMG in LLPS tendency. Despite being significant, the effect of mutation is moderate, likely because of the absence of a defined structure that may guide the selection of helix only weakening replacement.

Our results indicate that both N- and C-terminal domains are required for optimal LLPS, with the αMG structural element being crucial. The PTetC species has little tendency for LLPS and PNTet forms condensates but these are irregular, suggesting they are less fluid than those of full-length P. Based on these observations, an interaction network involving both P domains, namely, P_N-C_, P_N-N_ or P_C-C_ must form, and these partially overlap, i.e., protomers from either of the domain interact with at least two molecules in order for a network of weak interactions to be generated (Figure 8). In addition, we have to envision this network in three dimensions, so indeed different multiple combinations and arrangements are possible, much in the same way that both ionic and hydrophobic interactions participate. We hypothesize that the interactions are based on low specificity helix-helix interactions between different molecules and both domains, whose affinity is entropically favored from being tethered to the tight tetramerization domain that increases the effective concentration. In the case of the C-terminal domain, the helical components correspond to the dynamic αMG, and two transient helices that were described in the N-terminal domain. One of them corresponds to the binding site of N^0^ (the nucleoprotein without RNA) and the other to the transcription antiterminator M_2-1_. These proto-helices were shown to contact transiently and largely involve hydrophobic residues aligned at a face of the helix as previously indicated in wheel representations [53]. The irregular condensates formed by PNTet suggest evolution to more rigid material properties based on the hydrophobic interactions of the helices within. In the case of the C-terminal domain, the interprotomer helices that stabilize the αMG (α_C1_ and α_C2_) compete with interactions with N- or C-terminal helices from a different molecule (Figure 8A). Overall, the interactions stabilizing the α-helices in the N-term domain of P (α_N1_ and α_N2_) seems to be mainly hydrophobic while those at the C-term (α_C1_ and α_C2_) involve ionic interactions (Figure 8A, helical wheels, [53]. The C-terminal domain would thus provide fluidity to the condensates, and this is the reason for the formation of highly regular, reversible and dynamic droplets.

**Figure 8.**
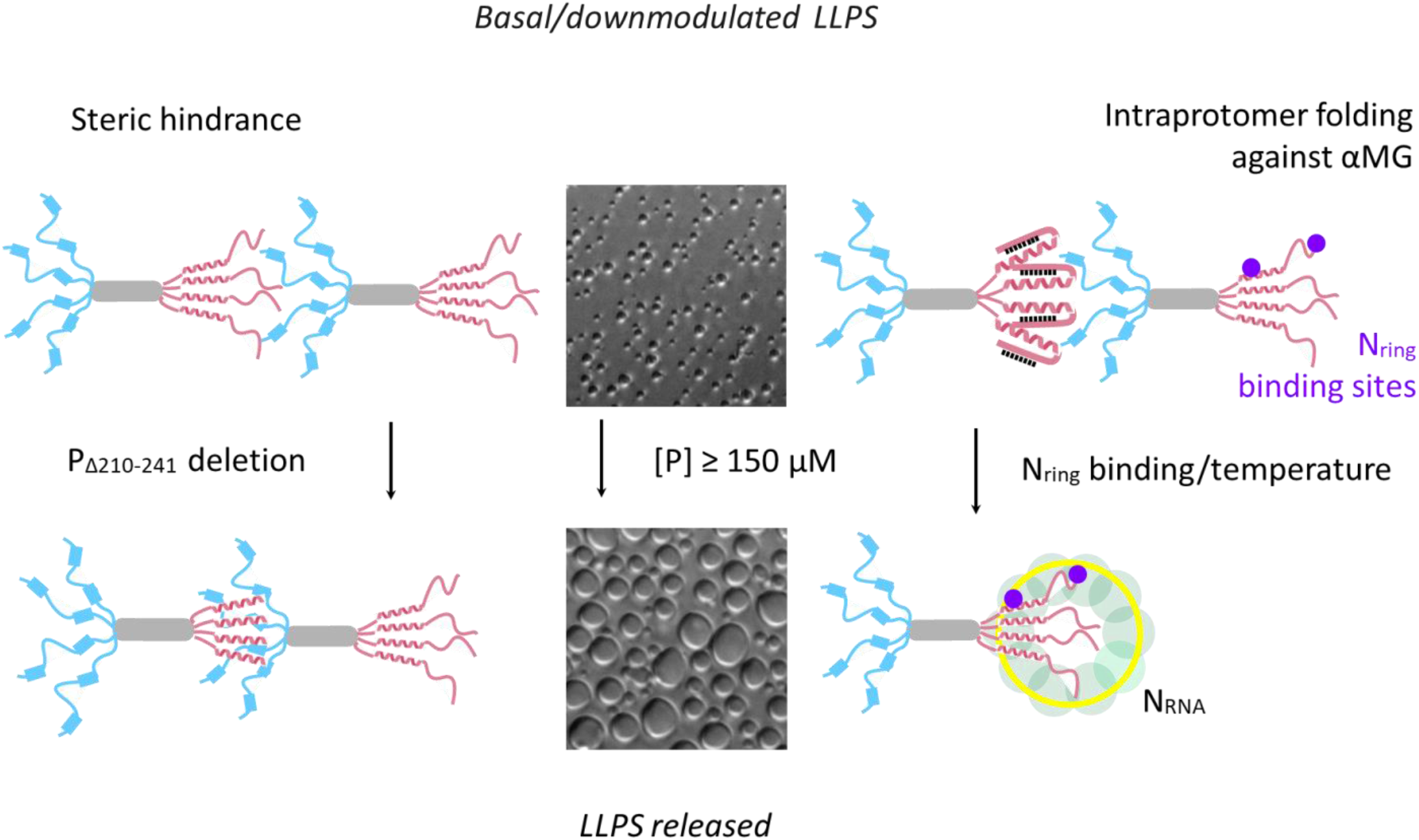
Hypothetical models for RSV-P LLPS modulation. Basal homotypic P LLPS is kept at a minimum in the full-length protein, which could be either by sterical hindrance between interacting domains (P_C-N_, P_C-C_, P_N-N_), which is released by deletion of residues from 210 to 241, or by binding of N_RNA_ to the reported binding site at the very C-terminus of P. Alternatively, the adjacent sequences could be involved in structured conformations folded against the αMG, which are released by temperature increase, binding of N_RNA_, or both. Magenta coloured spheres indicate both the primary known N_RNA_ binding site and a secondary binding site at the C-terminal domain of P (see main text).

A remarkable finding in this work is the presence of strong negative modulatory elements for LLPS at the C-terminus of P. Deletion of 10 residues in P_Δ232_ takes the LLPS onset temperature from 33°C to 44°C, and to 55°C when 32 residues were deleted in P_Δ210_ (Figure 3C). The structural basis of this negative modulation can be either the formation of an intraprotomer structure where the deleted elements fold against the αMG, or that these regions hinder the intermolecular interactions required to form the network of interactions between either of the domains (Figure 8). It should also be noted that the effect is partial with the small deletion, suggesting two structural components behind this negative effect. Intriguingly, a negative modulatory N binding element was identified by deleting the region spanning residues 198-217 of the C-terminal domain, mapping negative modulation of N binding and of LLPS to a similar region [62]. In addition, the ionic strength effect on homotypic LLPS is maximum at 150 mM NaCl for full-length P and the α3 mutant, but the deletion mutants show a different behavior. The LLPS of P_Δ232_ is much more sensitive to salt, where four negative charges are removed. On the other extreme, P_Δ210_ is not affected by ionic strength (Figure 3D), suggesting a predominant role of hydrophobic interactions for this variant.

Heterotypic N_RNA_-P condensation takes place *in vitro* at submicromolar concentrations (Figure 5B and S6) in contrast with dissociation constants of ca 30 µM reported from ITC and NMR [53, 63]. The relative concentrations of both components define a droplet to dissolution transition in excess concentration of P and droplet to aggregates in excess of N_RNA_. This shows a fine stoichiometric modulation in particular by the highly soluble P, which is able to rescue N_RNA_ aggregates readily into condensates (Figure 5C). Although N_RNA_ binding was mapped to the last nine residues at the C-terminal of P [48], we found that PNTet can form heterotypic droplets with N_RNA_, and both P_Δ210_ and P_Δ232_ deletions co-precipitate with N_RNA_, in all cases implying interactions in the absence of the binding site earlier described. In agreement with this, NMR line broadening of P in the presence of N_RNA_ also suggested a secondary binding site [53]. Thus, heterotypic LLPS is stabilized by multiple interactions involving more than one region, strengthened by the fact that the tetramerization domain increases the effective concentration of otherwise weak interactions. Downmodulated homotypic P LLPS is also released by binding to N_RNA_, presumably breaking the same negative interactions than P_Δ210_ and P_Δ232_ deletions release (Figure 8).

Granules or so-called inclusion bodies were observed for RSV N-P in cells [29, 32], and also for HMPV [54] ggand RABV [56] which we show their liquid nature and, importantly, the sequential time dependent assembly of these liquid condensates, starting from a large number of small puncta to a discrete number of large condensates (Figure 7D). These condensates do not require the presence of the N-terminal domain of P likely because of the large concentration they can attain in the context of the cytosol, but lack of this domain would not be viable in the virus as it contains the binding sites for N_0_ and M_2-1_, as well as other components. A similar time course sequential growth of granules containing P and N was observed in infection experiments, with large viral factories as the endpoint (Figure 7E). We propose that the formation of the viral factories is driven by N_RNA_-P condensation, starting with their interaction at early protein synthesis followed by nucleation of minimal droplets. These droplets grow to recruit M_2-1_ and L, which are synthesized at later stages generating an increase in transcription an therefore a subsequent increase in protein translation, in a positive feedback mechanism driven by condensation [9] (Figure 9). M_2-1_ will accumulate at the core of the factory condensate as previously shown, in structures named iBAGs (inclusion body-associated granules) [24, 64]. The vectorial nature of the position of the genes, conserved throughout non-segmented mononegavirales, leads not only to a sequential expression of the components, but also to different levels of the different proteins, with N and P being in large excess over M_2-1_ and L (Figure 9).

**Figure 9.**
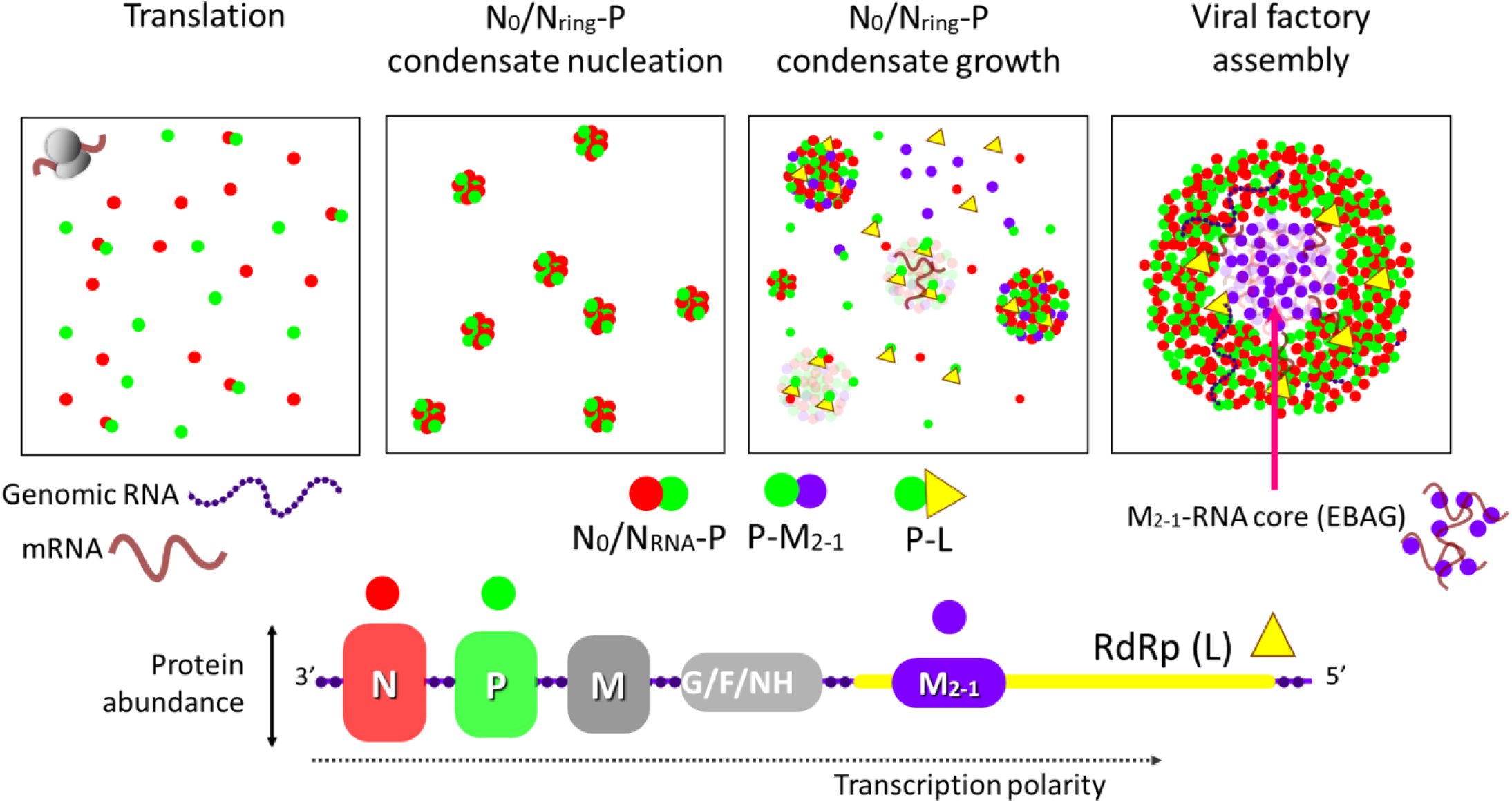
Biochemical mechanistic model for RSV viral factory assembly. See main text for explanation and also references [9, 24].

Clearly full-length P will not form homocondensates in the cell, as it will always be in the presence of the other viral proteins. However, understanding its mechanism of LLPS is essential since its scaffold nature means it will be driving condensation processes and given that, it is the only protein that interacts with the other viral replication proteins. Consequently, viral condensation will involve multiple condensation equilibria governed by P, in particular at the high concentrations attained in the viral factories. We propose that the condensation is driven by intermolecular interactions between weak transient α-helices of N- and C-terminal domains with those at the latter having a main role within a molten globule structure. These interactions are of low sequence/structure specificity, but are stabilized by the fact that are tethered to a tight tetramerization domain that allows for high effective concentrations allowing multiple interactions which would not take place in monomeric species.

The moderate LLPS tendency of full-length P strongly suggest that the appears to have evolved to be downmodulated at threshold levels that will undergo phase transitions with the subtlest changes, such as viral protein levels, changes in ionic strength, post-translational modifications, or binding to N. Its ability to interact weakly and strongly with viral proteins, its dynamic nature and capacity to stoichiometrically modulate the fluidity of condensates strongly suggest a role as the “solvent protein” within the viral factories, likely involving cellular proteins as well as viral mRNA and genomes. Future directions call for uncovering the nature of interactions and the role of water within the viral condensate in addition to those of high affinity already described, providing grounds for the exchange between dense phase (viral factories) and diluted phase (cytosol) and ultimately the transition to defined genome packages that will make up mature virions. All these stages defining the condensation mechanism provide a new and complementary platform for antiviral drug development [65], especially considering that it is a common mechanism in viruses, particularly RNA viruses, the likely candidates for emerging pandemics.

## MATERIALS and METHODS

### Protein cloning, expression and purification

All P sequences belong to human RSV A strain (optimized for *E. coli* expression). P and PTetC sequences were obtained by BamHI/EcoRI double digestion from a pRSET-P plasmid [41] and a pRSET-PTrI plasmid [51], respectively. PNTet, P_Δ210_ and P_Δ232_ sequences were amplified by PCR using pRSET-P as template. P and PTetC sequences were individually cloned into the BamHI/EcoRI sites of a pMal vector as N-terminal MBP fusion proteins, which contains an LVPRGS site for thrombin-mediated cleavage. PNTet, P_Δ210_ and P_Δ232_ were individually cloned into the BamHI/EcoRI sites of the pMal-C5X vector as N-terminal MBP fusion proteins, which contains an ENLYFQS site for TEV-mediated cleavage. α3mut has three mutations: G172S, R174A and E175G. The pMal-α3mut plasmid was obtained by site-directed mutagenesis using pMal-P as template. All resulting plasmids were sequenced and transformed in *E. coli* C41(DE3) for expression. A single colony was grown overnight at 37°C on 5 ml LB culture medium containing 100 µg/ml of ampicillin. The saturated culture was then used to inoculate 0.5 L LB medium containing 100 µg/ml. 0.4 mM IPTG was added at OD=0.6-1 for overnight induction. Cells were harvested by centrifugation at 4°C, resuspended in 25 mL of lysis buffer (100 mM Tris pH 8.0, 0.5 M NaCl, 2 mM β-mercaptoethanol, 1 mM EDTA and 1 mM PMSF), lysed by sonication and centrifuged at 9000 rpm for 40 min at 4°C. The soluble fraction was precipitated for 1 hour at 10°C and constant agitation by adding solid ammonium sulfate to a final concentration of 60% (w/v). The precipitated protein was collected by centrifugation at 9000 rpm for 20 min at 4°C, resuspended in 12.5 ml dialysis buffer (20 mM Tris pH 8, 200 mM NaCl) and dialyzed overnight against 4 L. The sample was subjected to affinity chromatography using amylose resin and eluted with elution buffer (20 mM Tris; pH 8.0), 200 mM NaCl, 20 mM maltose). The fusion protein concentration was determined by absorbance at 280 nm using the following molar extinction coefficients: 69330 M-1cm-1 for MBP-PTetC and 72310 M-1cm-1 for all other fusions. MBP-P and MBP-α3mut were cleaved overnight at 4°C using 0.6 units of thrombin per mg of fusion protein. MBP-PTetC was cleaved overnight at 4°C using 0.1 U/mg. The cleavage reaction was carried out in elution buffer supplemented with 2.5 mM CaCl_2_. MBP-PNTet, MBP-210 and MBP-232 were cleaved overnight at 4°C using a TEV:protein mass relationship of 1:300. After cleavage, all P variants were partially separated from MBP by reversible precipitation of P after addition of 4 volumes of 100 mM sodium acetate pH 4.6, 200 mM NaCl and incubation for 2 hours at 4°C. Precipitated protein was collected by centrifugation at 9000 rpm for 20 min at 4°C and resuspended in 100 mM Tris pH 8, 200 mM NaCl. The resulting resuspended protein was further centrifugated for 20 min at 4°C and 9000 rpm and subjected to size exchange chromatography using a Superdex 200 column, at a flow rate of 2 ml/min. P fractions were pooled, dialyzed overnight against 4 L stock buffer (10 mM Tris pH 8, 25 mM NaCl) and concentrated using centrifugal filter units to a concentration equal or higher than 150 µM. Protein concentrations of the P variants were determined by absorbance at 280 nm using the following molar extinction coefficients: P WT (6960 M-1cm-1), PTetC (3230 M-1cm-1), PNTet (6835 M^-1^cm^-1^), P triple (6835 M^-1^cm^-1^), P_Δ210_ (6835 M^-1^cm^-1^), P_Δ232_ (6835 M^-1^cm^-1^). N_RNA_ expression and purification protocol was described previously [38], although some minor modifications were made here. Briefly, a single colony of E. coli C41(DE3) transformed with the plasmid pRSET-N was grown at 37°C in 0.5 L LB medium containing 100 µg/ml amipicilin. At an OD=0.4, 0.4 mM IPTG was added for overnight induction at 22°C, yielding N-terminal 6xHis tagged N. The cell pellet was resuspended in 50 mM sodium phosphate (pH 8.0), 0.3 M NaCl, 2 mM β-mercaptoethanol, 1 mM EDTA and lysed by sonication. The soluble fraction was precipitated for 1 hour at 10°C and constant agitation by addition of solid ammonium sulfate to a final concentration of 60% (w/v). After resuspending and dialyzing against 20 mM sodium phosphate (pH 8.0), 0.3 M NaCl, the 6xHis-N was purified by standard Ni^2+^ chromatography and eluted with an imidazol gradient from 0 to 0.5 M in 30 minutes. The eluted fractions were pooled, dialyzed against 20 mM sodium phosphate (pH 8.0), 0.3 M NaCl overnight, concentrated and subjected to size-exclusion chromatography on a Superdex 200 column (Cytiva). Fractions were collected, pooled, further concentrated up to 150-200 µM. P is a tight tetramer with an unfolding/dissociation temperature above 80°C [41] and N_RNA_ rings are highly stable decameric structures [38], and therefore we express their concentration as tetramer and decamer, respectively, as they remain as such in all conditions.

### Use of crowder

Homotypic LLPS of full-length P was evaluated in 100 mM Tris-HCl (pH 8.0), 150 mM NaCl, and the optimal crowder concentration used was 15% PEG 4000 and a fixed FITC-P concentration of 0.1-0.5 µM. In the case of PtetC, the PEG concentration was increased to 25% to show the weaker tendency. Whereas heterotypic N_RNA_ and P phase separation was evaluated in NP-Ref buffer containing 100 mM Tris-HCl (pH 8.0), 150 mM NaCl, and fixed Cy5-N_RNA_ and FITC-P concentrations of 0.1-0.2 µM and 0.1-0.5 µM, respectively. Although >LLPS is observable in the absence of crowder, we observed a better reproducibility using 5% PEG. Homotypic LLPS was triggered by addition of highly concentrated protein or NaCl as the final mixture component. Heterotypic LLPS was triggered by addition of N or PEG 4000.

### Fluorescent labelling of proteins

All P species were labelled with Fluorescein isothiocyanate (Sigma-Aldrich). N_RNA_ was labelled with Cyanine5 NHS ester (Lumiprobe). All reactions were carried out overnight at 4°C, in sodium phosphate buffer (pH 7.0), 200 mM NaCl and at low dye:protein molar ratios (not higher than 2:1) to favor reaction to the protein N-terminus and thus minimize unwanted covalent modification of amino-containing residues. Simultaneous buffer exchange and purification from free dye was done in PD10 (GE) size exclusion column. Cy5-N_RNA_ aliquots were stocked in 20 mM sodium phosphate buffer (pH 8.0), 0.3 M NaCl. All FITC-P species were stocked in 10 mM Tris-HCl buffer (pH 8.0), 25 mM NaCl. Labelling ratio (dye/protein) was lower than 1 in every case, except for PNTet which was higher (Ratio = 1.2).

### Microscopy Imaging

Imaging was performed on a Zeiss Axio Observer 3 inverted microscope using 20X and 40X objectives for bright field (Hg Lamp) and epifluorescence (Colibri 5 LED illumination system). Samples were prepared at room temperature and analyzed at 22°C after incubation at the same temperature if required. Depending on experimental requirements, samples were either placed between two coverslips, or loaded into 96-well plates (Corning non-binding surface) and imaged at the bottom of the plate after droplet sedimentation. The set up for each experiment is indicated in the corresponding figure legend. Qualitative droplet dynamics was assessed by adding labelled protein to unlabeled pre-demixed samples and thoroughly mixing by pipetting in a microcentrifuge tube before placing the sample between coverslips.

### Fluorescence recovery after photobleaching

FRAP analysis was performed on a Zeiss LSM 880 Airyscan confocal laser scanning microscope with a Plan-Apochromat 63/1.4 objective lens at 22°C. 200 µL of 10 µM P (0.25 µM FITC-P) in reference condition was loaded into a Nunc Lab-Tek Chambered Coverglass (ThermoFisher Scientific Inc) and incubated for 2 h at 22°C. A circular region of interest (ROI) of droplets settled at the bottom of the coverglass were bleached using 90% laser power (n=6). A z-stack of three images were taken every 3 seconds, recording fluorescence intensity for three different ROIs (bleached droplet, reference droplet and background) over 210 frames, including 10 frames before bleaching. The fluorescence intensities were corrected as described elsewhere [66]. FRAP data was analyzed using the software FIJI Image J. FRAP curves were fitted to a bi-exponential function.

### NMR

For ^15^N-labeling, *E. coli* C41 (DE3) cells harboring the (PTetC) expression plasmid were grown at 37°C in M9 medium containing 1 g/L ^15^NH_4_Cl, and 100 mg/mL ampicillin up to an OD_600_ nm of 0.5. Recombinant protein expression and purification was performed as described above. NMR spectra were collected at 298 K on a Bruker Avance III 600 MHz spectrometer. Chemical shifts for PTetC were already available [53] and confirmed by ^15^N-TOCSY-HSQC and ^15^N-NOESY-HSQC. NMR data were processed with NMRPipe [67] and analyzed with NMRView [68] programs on a Linux workstation. Spectra were zero filled to 4 K and 0.5 K points in the t2 and t1 dimensions, respectively, and were apodized using a phase-shifted Gaussian function. The ^1^H and ^15^N resonance variations were followed at 288 K by collecting HSQC experiments. Chemical shifts perturbations (CSPs) were calculated from the spectra obtained in the absence and presence of 15% TFE using the following equation:

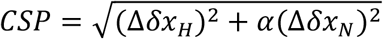

where Δδ*x*_H_ and Δδ*x*_N_ corresponds to the difference between chemical shifts in ^1^H and ^15^N for PTetC, and α is the scaling factor, it takes the value of α=0.2 for Gly and 0.14 for the rest of the residues.

### Far UV circular dichroism

All measurements were registered on a Jasco J-810 spectropolarimeter using a Peltier module for temperature control and a cell path length of 0.1 cm. CD spectra were recorded at standard sensitivity, a scanning speed of 100 nm/min, a time response of 4 seconds, a data pitch of 0.2 nm and a bandwidth of 4 nm. All spectra are an average of four consecutive scans.

For thermal scans, a single wavelength was recorded at standard sensitivity, a time response of 4 s, a bandwidth of 4 nm and temperature change rates of 1, 2 or 5°C/min. Data was collected every 0.5 or 1°C.

### Turbidity experiments

Since P and its variants are very stable and not prone to form irregular aggregates on the conditions that were analyzed, turbidity was used as quantitative reporter of homotypic LLPS. It was measured at 340 or 370 nm in either a spectrophotometer, a spectropolarimeter or a microplate reader depending on the requirements of the experiment. Homotypic LLPS thermal reversibility experiments and critical concentration determination were performed in a Jasco Spectrophotometer coupled to a water bath. Measurements were produced on time course mode, at 340 or 370 nm, and data was collected every 5 seconds. For thermal reversibility experiment, turbidity was registered at 340 nm in a 120 µl cuvette with a cell path length of 1 cm. A baseline of reference buffer was recorded at 10°C for 100 seconds, then P was added at a final concentration of 2.5 µM, thoroughly mixed by pipetting and the water bath temperature was set to 45°C. After achieving a new baseline, the temperature was reset to 10°C.

Critical concentration was determined by a dilution approach, monitoring scattering at 370 nm in a 700 µl cuvette with a cell path length of 1 cm at 15°C. A baseline of non-demixing condition was registered by measuring the samples at 30 µM protein in reference buffer without NaCl. Then LLPS was triggered by addition of NaCl at a final concentration of 150 mM and turbidity signal was recorded after incubation for 200 s. Dilution proceeded by taking a volume (100-350 µl) from the sample and adding the same volume of buffer 100 mM Tris-HCl, 15% PEG 4000, 150 mM NaCl without protein, preincubated at 15°C. TFE induction of homotypic demixing was determined by monitoring scattering at 370 nm in a 250 µL cuvette with a cell path length of 0.1 cm at 25°C. A baseline of 2.5 µM P containing 0, 5 or 10% TFE (v/v) was registered for 50 seconds and then NaCl was added to a final concentration of 150 mM and thoroughly mixed. For turbidity measurements in microplate format, 100 µL samples were placed in a 96-well microplate (Corning non-binding surface), incubated at room temperature for 0.5-1 h and measured in a Beckman Coulter DTX 880 Multimode Detector at 370 nm and 20°C unless stated otherwise. Determination of LLPS threshold temperatures of P variants was performed in a Jasco J-810 spectropolarimeter coupled to a Peltier module for temperature control, using a 250 µl cuvette with a cell path length of 0.1 cm, a response time of 4 s and a bandwidth of 4 nm. A baseline of reference buffer was registered in Absorbance mode at 370 nm and 65°C, then protein was added at a final concentration of 5 µM, incubated for 5 min and the temperature was lowered to 5°C. The cooling process was performed at a temperature change rate of 1°C/min, whereas reversibility was evaluated at 5°C/min.

### Cell Imaging

A549 cells (human lung carcinoma, ATCC reference: CL-185) were grown in DMEM-F12K medium supplemented with 10% fetal bovine serum. For immunofluorescence experiments of transfected cells, 105 A549 cells were transfected using Jetprime (Polyplus) with 0.5 µg of eukariotic expression plasmids pcDNA-P, -N, -PTetC, -PNTet, and/or -P fused with GFP. For immunofluorescence experiments of infected cells, a suspension of RSV wild-type strain A2 (106 PFU/ml) was used to infect A549 cell monolayer cultures at a MOI of 1. Infectious virus was adsorbed to the cells for 1 h in a 37°C incubator in 5% CO2. Following adsorption, the inoculum was removed and fresh medium added. After 24hs, cells were fixed with 4% paraformaldehyde, permeabilized for 30 min with 0.1% triton-X 100 and blocked with 3% BSA in PBS. Cells were then incubated overnight at 4°C with the indicated primary antibody (mouse IgG anti-P RSV or anti-N RSV), washed and incubated for 1 h with goat-anti mouse Cy3 conjugated IgG secondary antibody (Jackson). Images were obtained on a Zeiss LSM510 confocal microscope using a Plan-Apochromat 63x/1.4 Oil immersion objective.

For live-cell imaging, 2 104 A549 cells were seeded onto Nunc Lab-Tek chambered coverglass (ThermoFisher Scientific Inc). After 24hs, cells were co-transfected with pCDNA-GFP.P and pCDNA-N using Jetprime. Live-cell timelapse fluorescence was monitored at 488nm on a Zeiss LSM 880 Airyscan confocal laser scanning microscope (63×/1.4 oil-immersion objective) at 37°C and 5% CO2 over 24hs. For FRAP experiments, A549 cells were co-transfected as for live-cell experiments with pCDNA-GFP.P and pCDNA-N and analysed on a Zeiss LSM 880 Airyscan confocal laser scanning microscope with a Plan-Apochromat 63/1.4 objective lens at 37°C/5% CO2. 2.8 granule was bleached over 40 msec and recovery was monitored over 5 min.

## Notes

### Competing Interest Statement

The authors have declared no competing interest.

