## supplemental material for "Molten globule driven and self-downmodulated phase separation of a viral factory scaffold"

### SUPPLEMENTARY FIGURES AND TABLES

| [NaCl] (M) | [Diluted phase] (µM) | [Droplet phase] (mM) | [Droplet phase] (mg/mL) | PC* |
|---|---|---|---|---|
| 0.1 | 12.1 | 5.2 | 141 | 426 |
| 0.2 | 6.7 | 7.1 | 195 | 1070 |
| 0.3 | 4.9 | 6.8 | 187 | 1407 |

**Table S1. Homotypic partition of RSV-P in diluted and droplet phase**. 100 µl of 12.5 µM P (10% FITC labelled) in reference LLPS buffer at different NaCl concentrations were incubated at 4°C for 30 min and centrifuged at the same temperature for 10 min. The condensed phase was diluted 1/200 in reference buffer without PEG 4000 for Bradford quantification by duplicate, and confirmed by FITC fluorescence. *Partition coefficient calculated as [Droplet phase]/[Diluted phase]

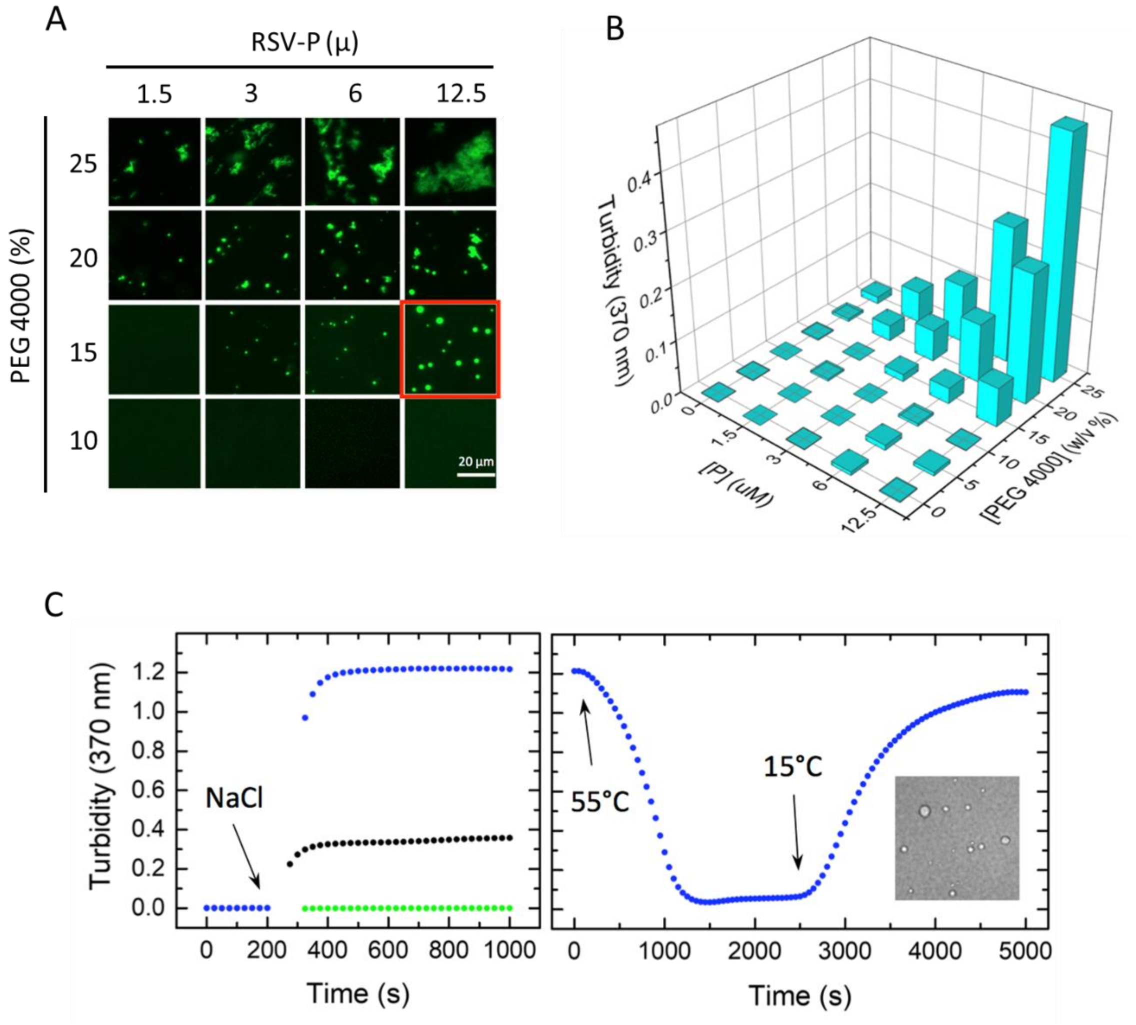


**Fig. S1. Effect of PEG 4000 on RSV P homotypic condensation.** RSV P condensation ability was tested by fluorescence microscopy (A) and turbidity (B). Samples containing increasing concentrations of both P and PEG 4000 in reference buffer were placed into a 96 well microplate, incubated for 1 h at 22°C and analyzed by turbidity at 370 nm and then by fluorescence microscopy at 40X. Microscopy images of samples containing less than 10% PEG 4000 and absence or protein are not shown. (C) Time course measurements of turbidity at 370 nm were taken at 15°C for samples containing 5 µM P WT (black dots), 25 µM PTetC (green dots), both in reference condition, or 25 µM PTetC in the presence of 25% PEG 4000 (blue dots). Demixing was triggered by addition of NaCl to a final concentration of 150 mM. The effect of temperature on PTetC demixing was tested by heating the sample to 55°C and then cooling back to initial temperature. Inset: PTetC droplets at 20°C at 100 µM and 25% PEG 4000.

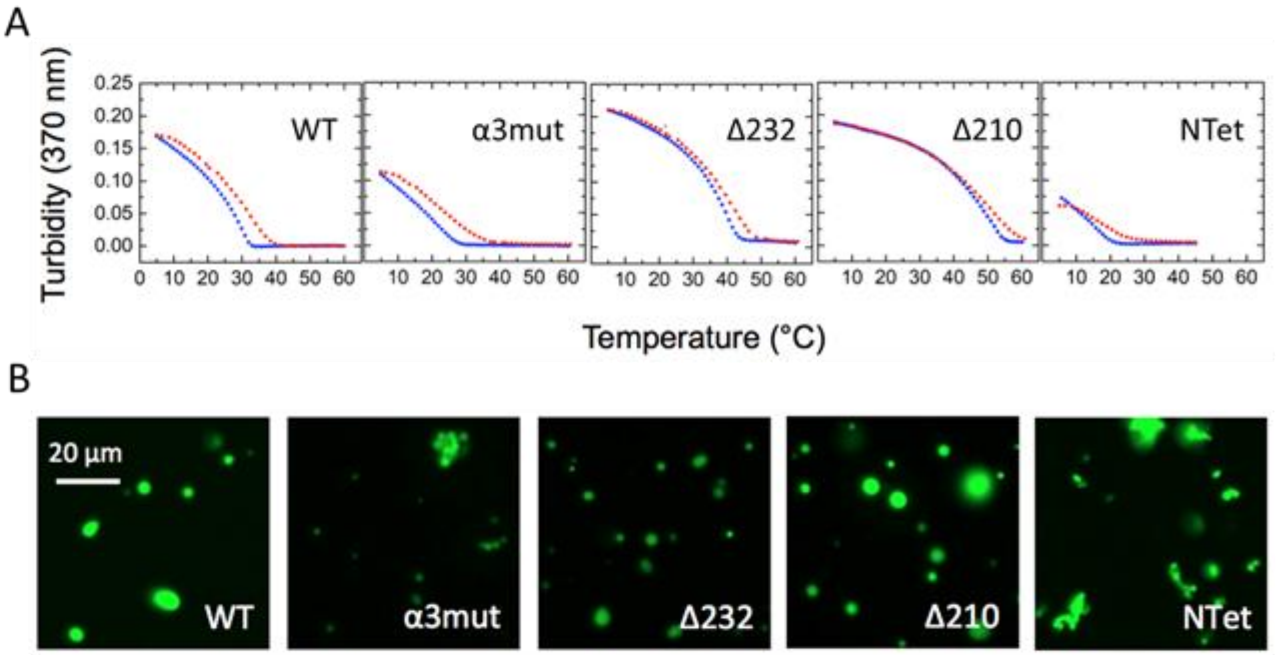


**Fig. S2. Droplet dynamics and reversibility of P and its variants.** (A) Reversibility of droplet formation for P WT and its variants was tested by thermal scans followed by turbidity at 370 nm. Cooling (blue dots) was performed at 1°C/min. Heating (red dots) was done at 5°C/min. **(B)** Fluorescence microscopy images of samples containing 12.5 µM protein in reference condition at 22°C. Images were acquired at 40X 5-10 min after addition of 0.25 µM FITC-labelled protein to non-fluorescent pre-demixed samples that were previously incubated for 1 h at 22°C.

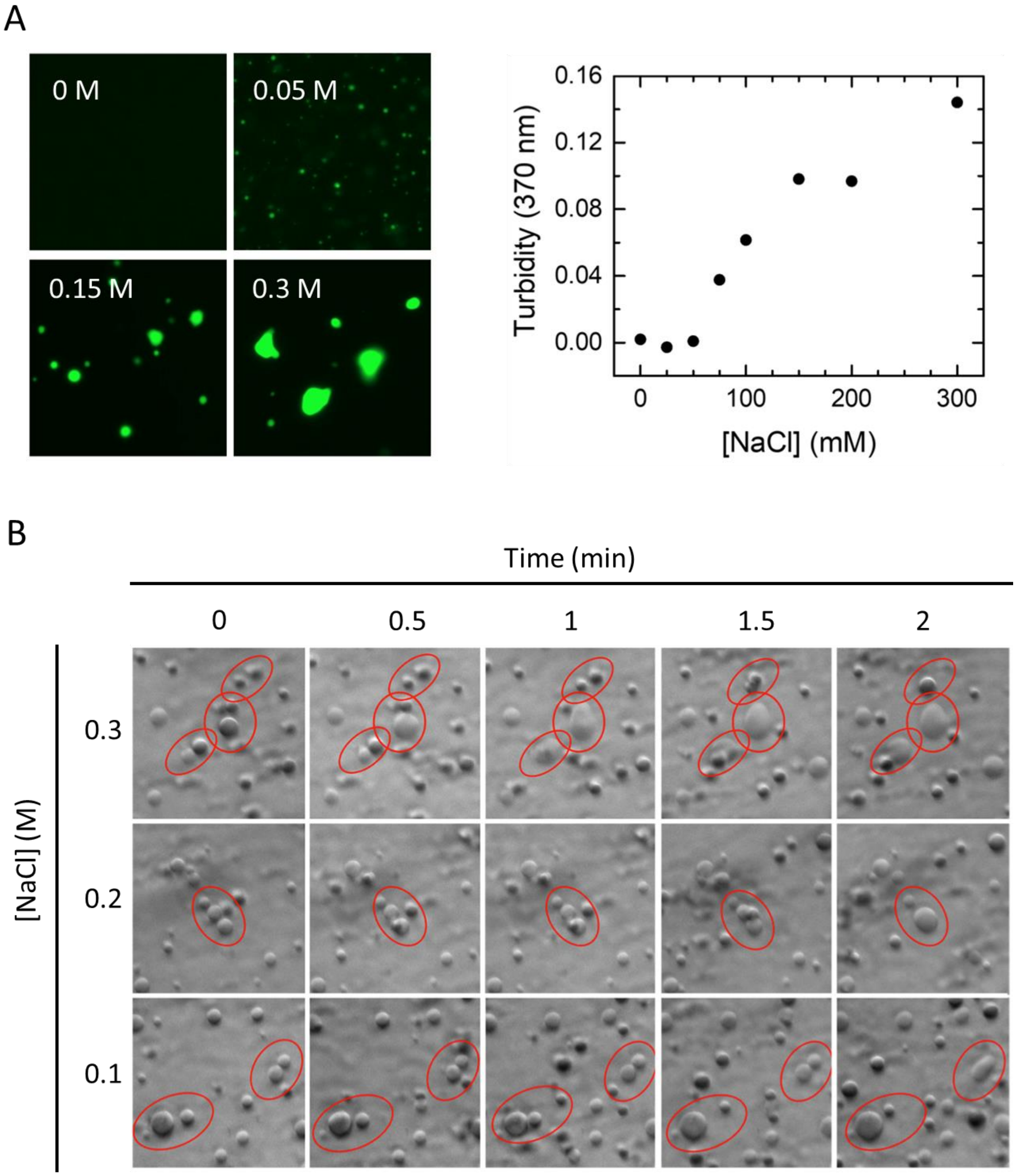


**Fig. S3. Effect of ionic strength on P homotypic LLPS**. (A) Fluorescence microscopy images (left) and turbidity (right) of samples containing 12.5 μM P WT in the presence of increasing concentrations of NaCl. (B) Coalescent events of P WT droplets at different NaCl concentrations registered by bright field microsocopy at the bottom of a 96-well microplate.

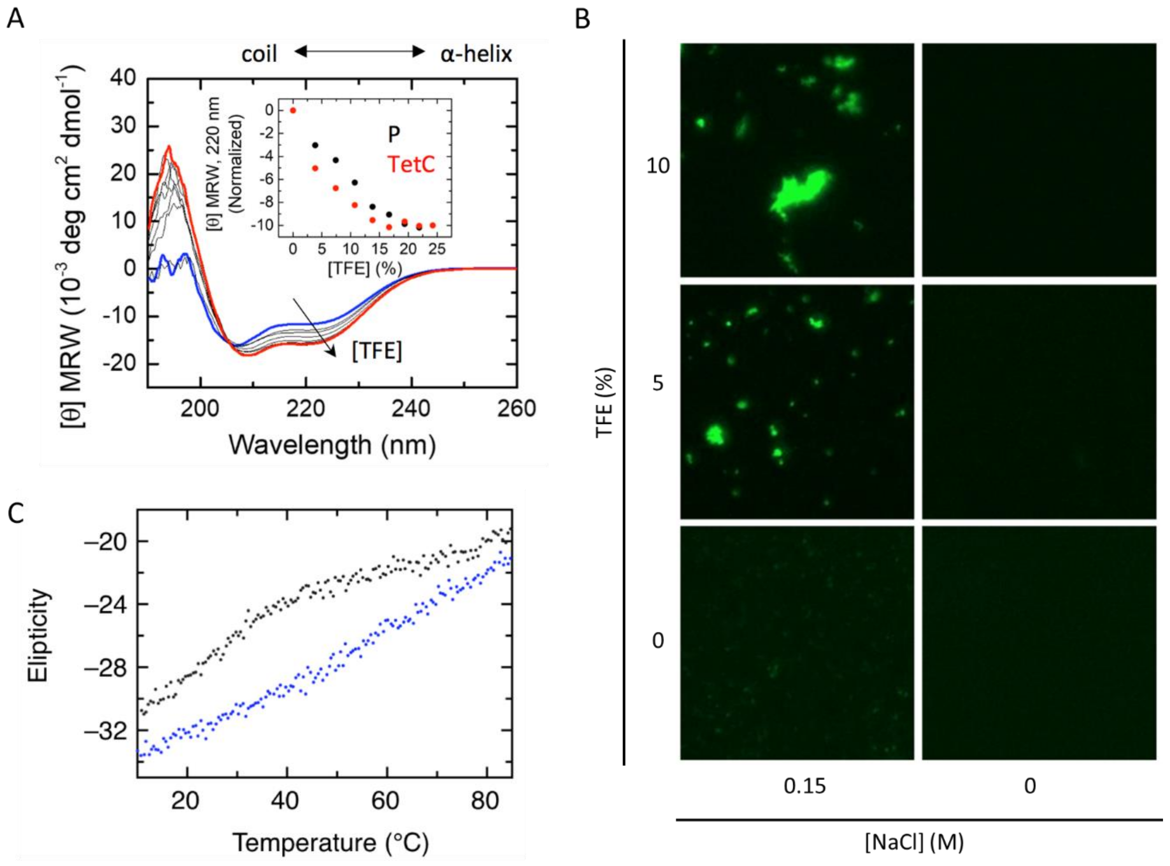


**Fig. S4. Stabilization of MG domain by TFE. (A)** TFE titration of P WT or PTetC were titrated with TFE up to a final concentration of ca 25% (v/v) and Far-UV CD spectra were recorded at each titration step after a three minute incubation at 15°C. Inset: normalized molar ellipticity of P WT and PTetC at 220 nm as a function of [TFE]. Samples contain 3.75 µM protein in 10 mM Tris-HCl (pH 6.5), 25 mM NaCl. **(B)** Effect of TFE on P WT condensation. Demixing of 2.5 µM P in reference buffer was evaluated by turbidity at 370 nm after addition of 0.15 M NaCl in the absence (black circles) and presence (red circles) of 5% and 10% TFE. Fluorescein-labelled P samples were then prepared in the absence and presence of 0.15 M NaCl and analyzed by fluorescence microscopy at 20°C. Scale bar = 20 µm. **(C)** Thermal scans followed by CD at 220 nm of P WT (black) and α3mut (blue).

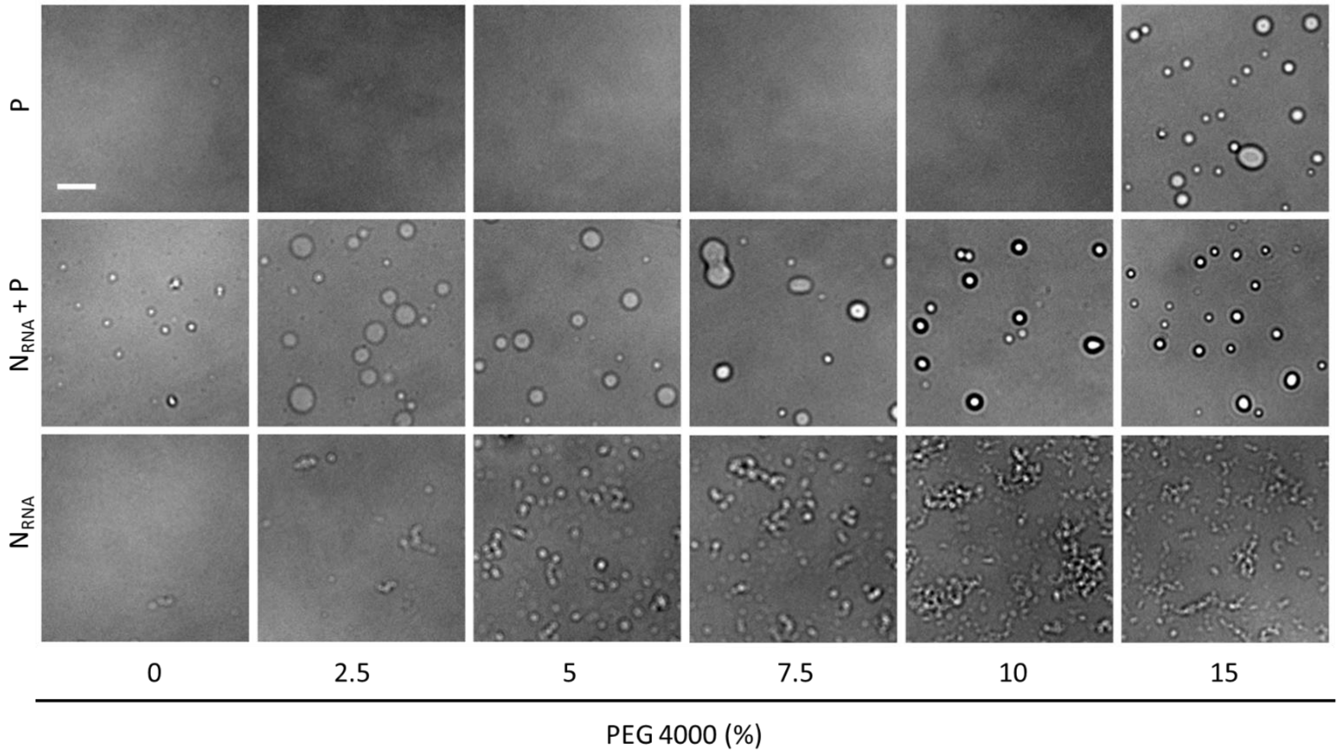


**Fig. S5. Effect of crowding on $N_{RNA}$-P LLPS and $N_{RNA}$ alone.** Bright-field microscopy images of P, $N_{RNA}$ or P+$N_{RNA}$ in the presence of increasing concentrations of PEG 4000. Imaging was performed by loading 1 µL of sample between two coverslips. [P] = 12.5 µM; [$N_{RNA}$] = 1.25 µM. Distance bar = 20 µm.

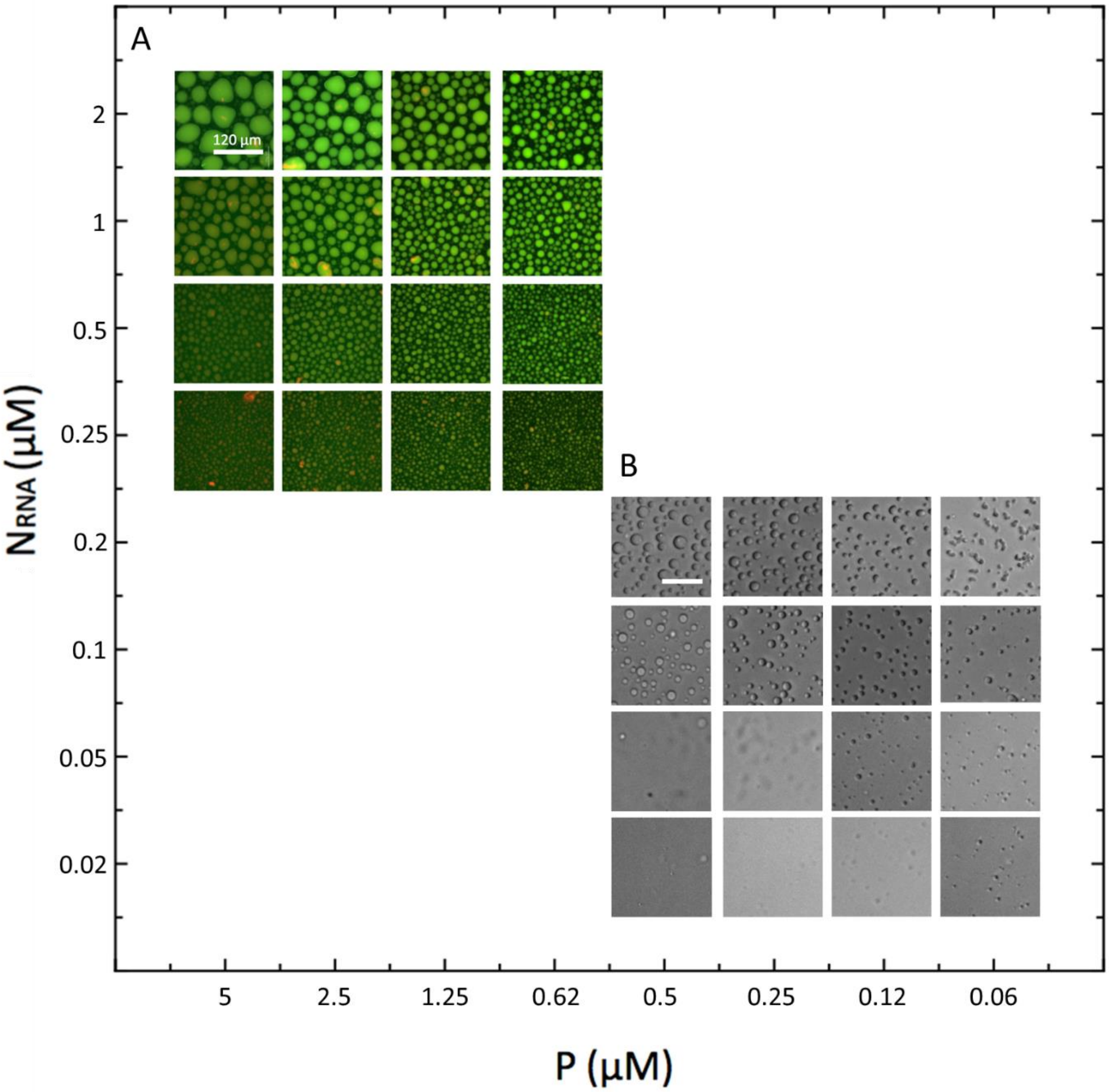


**Fig. S6. P : $N_{RNA}$ stoichiometry effect on heterotypic LLPS.** Two grids spanning different concentration ranges of $N_{RNA}$ and P in reference condition were analyzed by fluorescence **(A)** or bright-field **(B)** microscopy. All samples were loaded onto 96-well microplates and allowed to settle for 3 h at 22°C before imaging at the bottom of the plate.

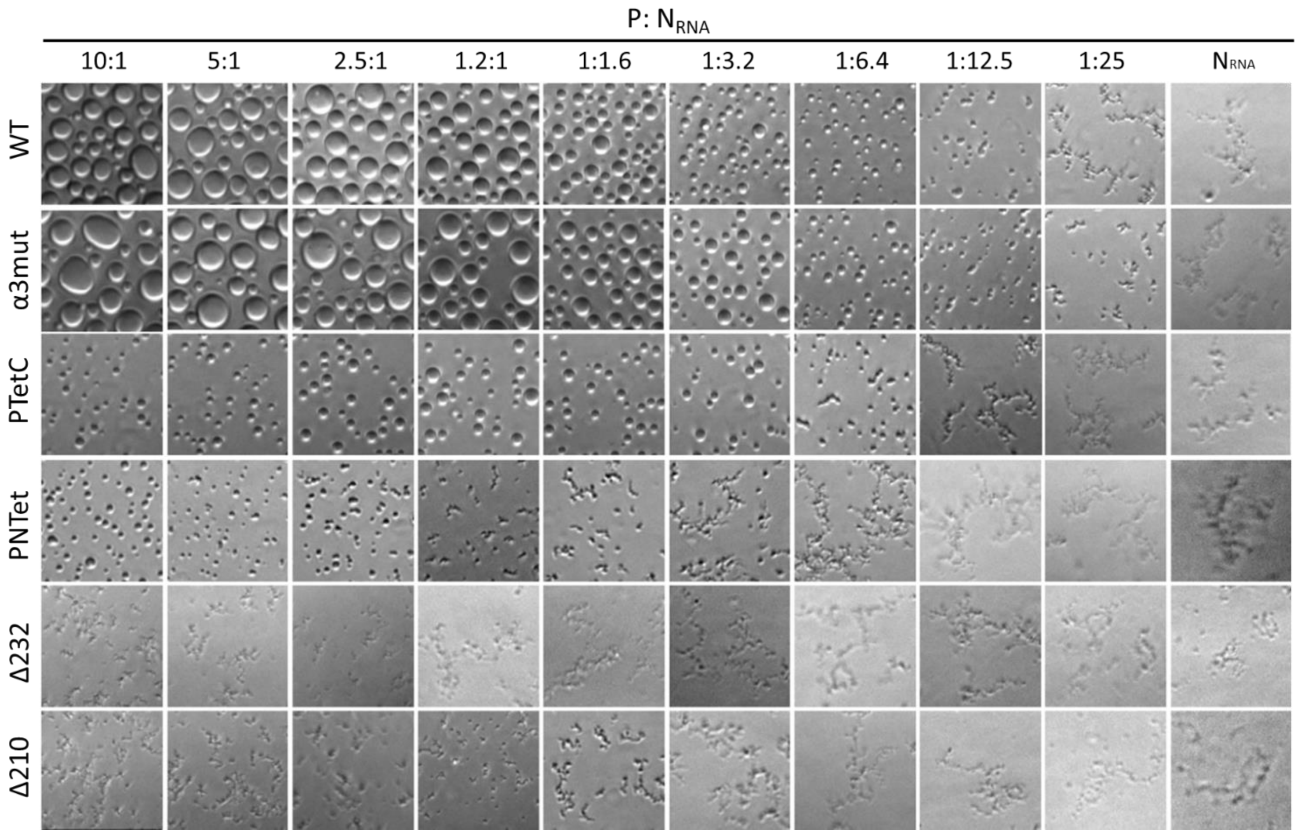


**Fig. S7. Analysis of P sequence regions involved in P:$N_{RNA}$ heterotypic LLPS.** Bright-field microscopy images of samples containing a fixed [$N_{RNA}$] = 0.5 µM and different concentrations of P variants spanning a wide stoichiometry as indicated in the upper side of the figure. All samples were loaded onto a 96-well microplate and incubated for 3 h at 22°C before imaging.

**Movie S1.**

MovieS1.avi

**P-N$_{RNA}$ condensates assembly in A549 cells**. Formation of P-N condensates in A549 cells co-transfected with pcDNA-N and pCDNA-GFP.P plasmids over 15hs. The evolution of the granules was registered using a Zeiss LSM 880 Airyscan confocal laser-scanning microscope at 37°C and 5% $CO_2$.
